# Ligand binding remodels the oligomeric state and active site of *Klebsiella pneumoniae* azoreductase

**DOI:** 10.64898/2026.09.29.755508

**Authors:** Reese Hitchings, Agnidipta Ghosh, Arthur J. Arcinas, Steven C. Almo, Libusha Kelly, Ali Ryan

## Abstract

Azo dyes are widespread in the environment, and humans ingest a range of these dyes present in food and drugs. The human gut microbiome significantly impacts the fate of ingested azo dyes primarily through the reduction of the azo bond (R−N=N−R′). Flavin-dependent azoreductases have been considered the key enzymatic drivers of this process. To challenge this paradigm, we constructed a sequence similarity network of known azoreductases and focused on AzoA of *Klebsiella pneumoniae*, a central representative within the gut microbiome cluster. Unexpectedly, *K. pneumoniae* AzoA displayed relatively low *in vitro* activity against a range of azo compounds. While the physiological implications remain unclear, these findings are consistent with the possibility that non-enzymatic mechanisms contribute substantially to azo compound reduction in the gut. Furthermore, structural characterization of AzoA, as a holoenzyme and in its substrate-bound state, revealed a previously unrecognized ligand-coupled mechanism involving remodeling of the active site and oligomeric state.

## INTRODUCTION

Azo compounds, characterized by the R-N=N-R’ azo bond, are among the most widely used synthetic molecules, constituting over 60% of all commercial dyes.^1^ Their applications are extensive, particularly within the food and pharmaceutical industries. The public is frequently exposed to azo compounds, as they include three of the four most common artificial food colorings (AFCs), such as Red 40, sunset yellow and tartrazine.^2, 3, 4^ Children in the U.S. alone can ingest over 60 mg of AFCs daily through processed foods and beverages.^5^ Azo compounds also have a long history in medicine, dating back to the first clinically available antibiotic, prontosil.^6^ Today, two main groups of azo drugs remain in clinical use, including those for inflammatory conditions like mesalamine and agents for urinary tract infections like phenazopyridine.^7, 8^

The therapeutic action of early azo drugs, like prontosil, was found to rely on its function as a prodrug; the active molecule, sulphanilamide, is released only after the enzymatic reductive cleavage of the azo bond.^9^ This cleavage, however, is not exclusive to drug activation. It is a common metabolic pathway for all ingested azo compounds and often results in the production of potentially toxic aromatic amines.^10^ Landmark studies in the 1960s demonstrated that the bacterial enzymes in the gut microbiome, the plethora of species of bacteria, archaea, and viruses inhabiting the human gut, are the primary drivers of this reductive azo cleavage. ^11–14^ This specific metabolic capacity has garnered significant attention for the targeted delivery of azo-containing prodrugs to the colon, offering promise for conditions like colorectal cancer and inflammatory bowel disease (IBD).^15, 16^

Crucially, overall azoreduction capacity in an individual can vary significantly,^17^ likely due to differences in the composition of their gut microbiome. This inter-individual variability has profound implications for both the toxicity of ingested azo dyes and the efficacy of azo prodrugs.

Despite the widespread human exposure to azo compounds and the critical role of the gut microbiome in their metabolism, little is known about the specific enzymes responsible for azo bond cleavage.^18^ Azoreductases, which are NAD(P)H-dependent flavoenzymes, are presumed to be the key players in this process.^18^ These enzymes belong to the NAD(P)H quinone oxidoreductase superfamily, which shares a common short-flavodoxin-like fold^18^ and are primarily homodimers in solution^19, 20^ with a few exceptions such as homotetrameric azoreductase, AzoR1 of *Pseudomonas aeruginosa*.^21–23^ The active site of azoreductases resides at the dimer interface involving residues from both protomers of the dimer. The non-covalently bound flavin cofactor engages a conserved network of sequence invariant hydrogen bonds involving mainchain carbonyl and amide groups in the L7 and L11 loops/segments within the flavodoxin-like fold.^24^ The sequence-independent nature of these interactions allows for significant variation in the amino acid sidechains surrounding the active site. The active site of the azoreductases is configured not to accommodate both the nicotinamide cofactor and the electron acceptor simultaneously, hence all enzymes in the family operate via a *bi-bi* ping pong mechanism.^23, 25^

Studies, however, have often shown that purified azoreductases exhibit unexpectedly poor activity against common azo dyes,^26, 27^ challenging the assumption that they are the sole or primary drivers of azo metabolism in the gut. To date, only a limited number of azoreductases from relevant gut bacteria have been characterized.^27^

To address this knowledge gap, this study characterizes a representative azoreductase from the human gut microbiome. Using a sequence similarity network (SSN) of all annotated azoreductases in the NCBI database, we selected AzoA (UniProt ID A6TCS9) of *Klebsiella pneumoniae*, a central enzyme within the largest cluster (cluster 1; Figure 1) of gut-associated azoreductases and found in 6-23% of gut microbiomes.^28, 29^ Here, we report the biochemical properties of AzoA and identify a unique mechanism of substrate-driven activation, providing novel insights into the varied and complex enzymatic systems that govern the metabolism of azo compounds in the human gut.

**Figure 1.**
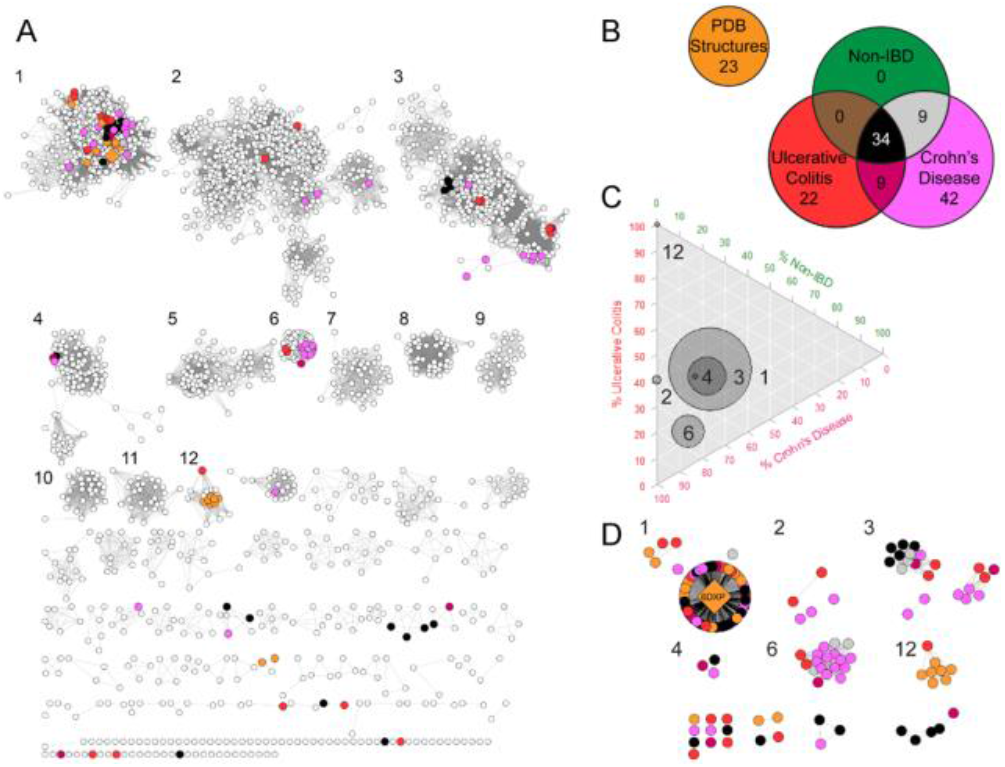
Global and human gut microbiome azoreductase sequence relationships and disease associations. (A) Sequence Similarity Network (SSN) depicting all unique azoreductase sequences from the NCBI protein database (BLASTP e-value ≥1*10^−20^). Individual clusters are numbered and nodes are color-coded to denote azoreductase sequences sourced from human gut DNA metagenomes within the IBDMDB. (B) A Venn diagram illustrating the IBDMDB disease metadata categories used for the coloring in panel A. (C) Ternary plot representing the distribution of patient disease metadata within each cluster of panel A. Circle diameter is proportional to the count of human microbiome-associated sequences per cluster. (D) A filtered SSN showing only the human gut metagenome azoreductases, maintaining the cluster identities and numbering established in panel A.

## RESULTS

### Azoreductase diversity analysis

To assess the diversity of the azoreductase superfamily, we retrieved 3,022 unique sequences from the NCBI protein database^30^ and applied 95% identity dereplication (Supplemental data 1). Of these, 116 sequences showed strong similarity to predicted proteins from the publicly available IBDMDB metagenome assemblies.^31^

We visualized the extensive variety using a Sequence Similarity Network (SSN) (Figure 1). This SSN maps each unique sequence as a node, with edges connecting sequences with a BLAST e-value <1×10^−20^, deemed “high-identity homologs”.

The resulting network demonstrates the extreme diversity of azoreductase-associated proteins: the 3,022 sequences segregate into 16 major clusters (each with ≥15 members; Figures 1 and 1B). Clusters 1 (1,182 sequences), 2 (666 sequences), and 3 (347 sequences) dominate the sequence space. Although automated text-based search includes many mis-annotated and irrelevant protein sequences, our approach successfully grouped and distinguished diverse enzyme families historically annotated as azoreductases, such as the FMN-dependent azoreductases (*E. coli* AzoR, Cluster 1; UniProt ID P41407),^32^ NAD(P)H oxidoreductases (Cluster 3),^33^ and arsenic resistance proteins (*P. aeruginosa* ArsH, Cluster 6).^34^ Despite their diversity, sequences from clusters 1, 3, 6, 11, and 12 have previously been reported to act as azoreductases (Figure S1 and Table S1).

### Elevated Arsenic Resistance Proteins in Crohn’s Disease

Our analysis indicates that arsenic resistance proteins are over-represented in patients with Crohn’s Disease (CD). Specifically, Cluster 6 of the azoreductase SSN is largely dominated by sequences from CD patients; only two out of 22 gut-associated sequences in this cluster originated from non-CD individuals (Figures 1C and 1D).

When these human gut-associated sequences were queried against the NCBI non-redundant protein database using BLASTp, they showed >90% identity to proteins annotated as arsenic resistance proteins. The top BLASTp hits were identified as ArsH proteins from diverse microbial species, including *Pseudomonas fluorescens, Citrobacter freundii*, and *Achromobacter xylosoxidans*.

These ArsH proteins share the short-flavodoxin-like fold characteristic of the enzymes in Cluster 1, and several are known to exhibit azoreductase activity.^34, 35^ While their name reflects their role in conferring resistance against the toxic metal arsenic and their critical part in the ecological arsenic cycle,^36^ ArsH from *Pseudomonas putida* has also been demonstrated to protect bacteria from oxidative stress.^37^

Though no causal link has been established between the presence of gut microbial ArsH proteins and CD, Cluster 6 is the only large cluster in the network that is overwhelmingly dominated by a single disease state. Together, these results suggest the presence of functionally-distinct and population-specific azoreductase subfamilies in the gut microbiome that may alter azo-compound metabolism.

To experimentally characterize azoreductases identified from the human gut microbiome, we synthesized 63 open reading frames (ORFs; Supplemental data 2) selected to represent the sequence diversity observed across the identified clusters and evaluated their heterologous expression. Of these, only nine ORFs yielded soluble recombinant proteins suitable for biochemical characterization (Figure S2). The successfully expressed proteins and their corresponding UniProt accession numbers are listed in Table S2. Notably, all soluble proteins originated from Cluster 1, despite the initial selection spanning multiple sequence clusters (Figure 1D). This finding highlights the current experimental bias toward Cluster 1 enzymes and underscores the challenges associated with expressing and characterizing more divergent members of this protein family using a standard aerobic *E. coli* expression system.

An azoreductase, AzoA, encoded by *Klebsiella pneumoniae*, was selected for subsequent enzymatic and structural studies based on two key criteria: **1**) It displayed the highest degree of centrality within Cluster 1 of the SSN, indicating it had the most connections to other sequences in that cluster; and **2**) It showed highest level of expression in *E. coli* (Experimental Methods).

### Biochemical characterization of recombinant AzoA

We expressed and purified recombinant hexa-histidine tagged AzoA from *Klebsiella pneumoniae* (Experimental Methods). Purification involved immobilized metal affinity chromatography (IMAC) followed by size exclusion chromatography (SEC; Figure S2A), yielding a highly pure protein (Figure 2A). The purified AzoA protein was bright yellow and showed characteristic absorbance peaks at 360 and 450 nm (Figure 2B). The ratio of absorbance at 450 nm (for the flavin cofactor) to 280 nm (for the protein) suggests that AzoA binds flavin mononucleotide (FMN) with a stoichiometry of nearly 1:1. To determine enzyme activity and cofactor selectivity, enzyme assays with AzoA were conducted under strict anaerobic conditions that mimic the gut environment.

**Figure 2.**
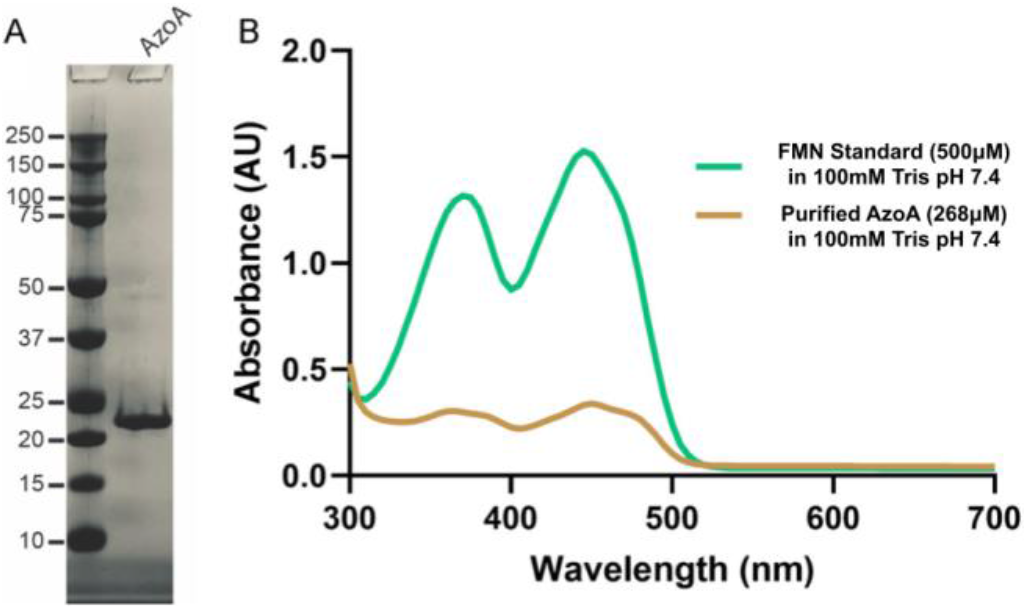
Basic characterization of recombinant AzoA of *K. pneumoniae*. (A) Aliquots (3 μg) of the peak Superdex S200 fraction of *K. pneumoniae* AzoA (Experimental Methods) were analyzed by SDS-PAGE. The positions and sizes (kDa) of marker proteins are indicated on the *left*. (B) UV-Vis absorption spectra of the AzoA and representative flavin cofactors as reference.

Initial experiments to gauge nicotinamide cofactor selectivity using menadione as the electron acceptor showed that AzoA does not have a significant preference for the electron donor; the rate of nicotinamide oxidation was comparable when using NADH or NADPH (Figure 3A). The addition of exogenous FMN caused a small but statistically significant increase in the overall reduction rate with both NADH and NADPH (Figure 3A).

**Figure 3.**
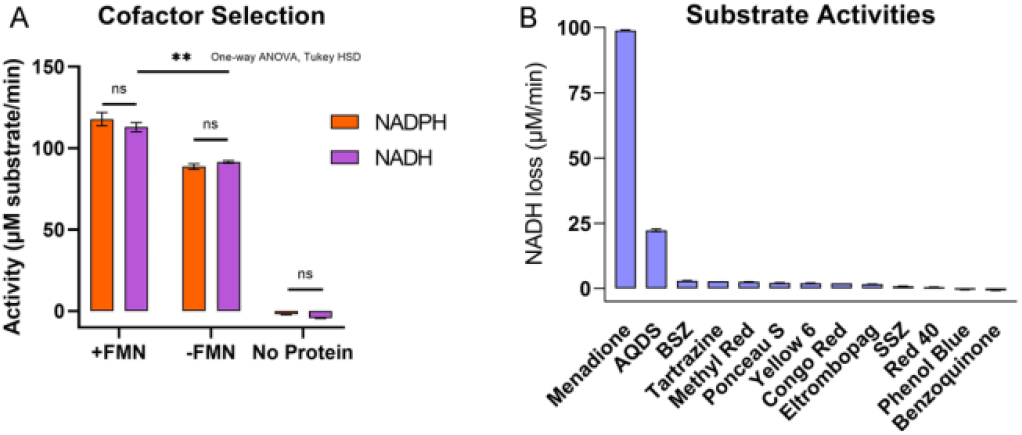
Enzymatic characterization of AzoA. (A) The nicotinamide cofactor selection of *K. pneumoniae* AzoA using menadione as the substrate. The rate of cofactor utilization was measured by monitoring the consumption of NADH and NADPH. The data reveal that the AzoA utilizes both NADH and NADPH at a similar rate. Furthermore, the rate of NADH consumption significantly increased upon the supplementation of flavin mononucleotide (FMN), as determined by a one-way ANOVA with Tukey HSD (p <0.01) (B) The activity screen of the AzoA against menadione, AQDS (organosulfonate), various azo and quinone substrates (Table S3). Enzyme activity was quantified by measuring the rate of NADH utilization, which was monitored spectrophotometrically at a wavelength of 340 nm. This analysis helps to identify which substrates the AzoA enzyme can effectively act upon.

We tested a range of substrates to determine AzoA’s selectivity, including food colorants (AFCs like tartrazine) and azo prodrugs (e.g., balsalazide) (Figure S3). The reduction rates for these traditional azo substrates were only barely detectable (Figure 3B; Table S2) under the assay condition tested (Experimental Methods). Given that azoreductases belong to the NAD(P)H quinone oxidoreductase family, we tested several soluble quinones. While benzoquinone resulted in NADPH oxidation rates similar to those observed with the azo dyes, rates with naphtho- and anthraquinones (such as menadione) were significantly faster.

### The structures of the tetrameric AzoA holoenzyme and dimeric Red 40 bound AzoA

To probe substrate selectivity and quaternary assembly, the structure of the AzoA holo-enzyme was determined, and the final model was refined to 2.48 Å with four protomers in the asymmetric unit (PDB 6DXP; final R_work_ and R_free_ values of 23.7% and 28.5 %, respectively). Crystallographic data statistics are summarized in Table S4. Each protomer of AzoA forms a short-flavodoxin-like fold and after initial phasing additional electron density was observed in the active sites of all four protomers allowing incorporation of FMN into the model. The four chains within the asymmetric unit form a tetramer composed of a dimer of dimers (Figure 4A) with a large total buried surface area of 5132 Å^2^ (Figure 4B). PISA analysis predicts that this tetrameric conformation is stable in solution (ΔG= −70.6 kcal/mol),^38^ which is consistent with data obtained from Dynamic Light Scattering (DLS; Figures 5 and S4). A critical feature of the structure of the AzoA holoenzyme is the positioning of the side chain of Gln145 close to the N10 atom of FMN (3.4 ±0.17 Å; n = 4) and shielding of the cofactor (Figures 4C). The position of Gln145 in all chains is stabilized via a short range (2.6 Å) hydrogen bond to Tyr120 sidechain hydroxyl (Figure 4C).

**Figure 4.**
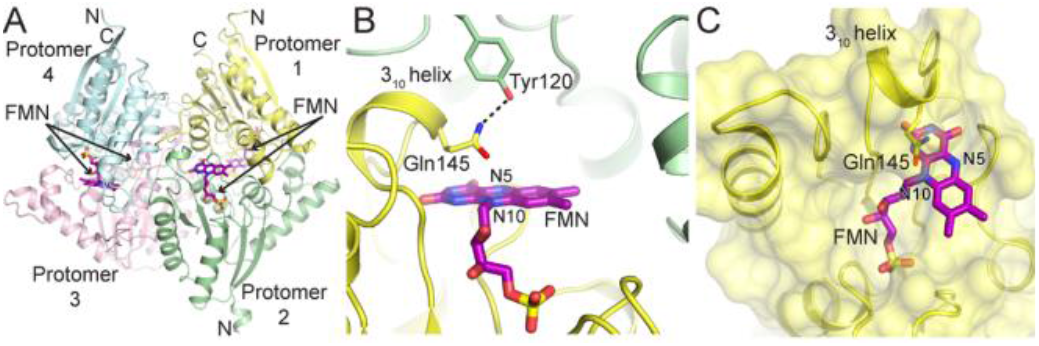
The Tetrameric Structure and Active Site of *K. pneumoniae* AzoA Holoenzyme. (A) The crystal structure of the AzoA tetramer a ribbon with arrows for β-strands and wide ribbons for helices. Four protomers (1-4) are color coded in yellow, green, light pink and cyan, respectively. Location of active site in each protomer is shown with FMN (stick representations; purple). N and C denote N- and C-termini. (B) A close-up view of an AzoA active site showing bound flavin mononucleotide (FMN) cofactor in purple stick representation. The panel highlights the 310 helix (part of the L11 segment), which contains the amino acid Gln145. A dashed line indicates the interaction between Gln145 and Tyr120 from an adjacent subunit (in *trans*). Atomic contacts are indicated by dashed lines. (C) A transparent surface representation of a single AzoA subunit shows how the side chain of Gln145 within the same subunit (in *cis*) acts as a shield, protecting the bound FMN from the solvent.

**Figure 5.**
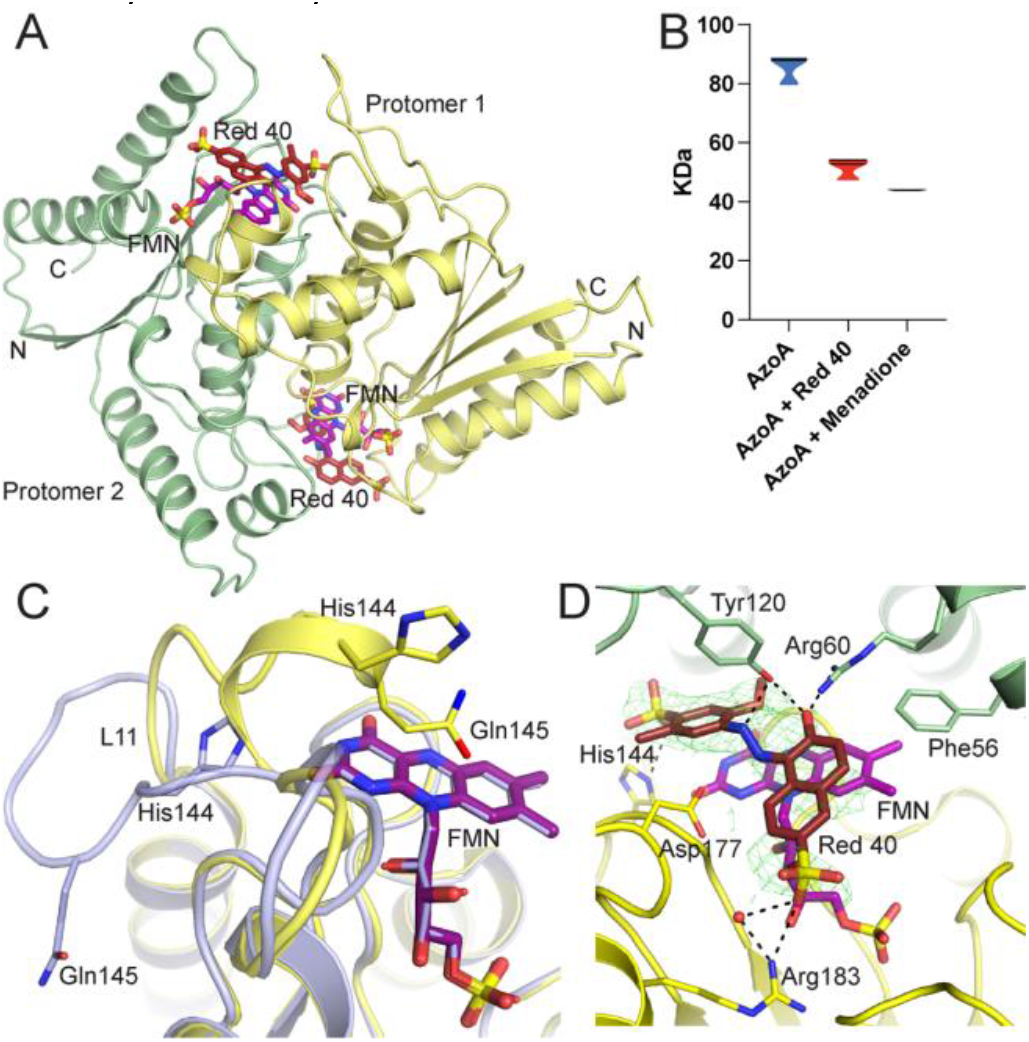
The structure of *K. pneumoniae* AzoA bound to Red 40. (A) Crystal structure of AzoA bound to its cofactor, FMN (purple), and the azo dye Red 40 (dark red sticks). The two AzoA subunits are represented as in Figure 4A. (B) Violin plots display the apparent molecular weight (MW) distribution of AzoA alone, with Red 40, or with menadione. The MW was calculated from dynamic light scattering (DLS; Figure S4) data, which measured the hydrodynamic radii of the samples (Experimental Methods). (C) Conformational changes in the L11 segment of AzoA upon binding Red 40. The structure of the apoenzyme (without Red 40; Figure 4C) is shown in yellow, while the Red 40-bound structure is in light blue. The FMN molecules for each subunit are shown in purple and light blue, respectively. Key amino acid side chains involved in the conformational change are shown as sticks and labeled. (D) Closeup view of the AzoA active site. The FMN cofactor is shown in purple, and the bound Red 40 is in dark red. A green mesh represents the electron density (omit map) of the bound Red 40, contoured at 2.5σ, confirming its presence in the active site. A water molecule interacting with Red 40 is shown as a red sphere. Atomic contacts are indicated by dashed lines.

To provide a structural benchmark for comparison, we also determined the crystal structure of the well-characterized *E. coli* azoreductase AzoR, which shares ~50% sequence identity with *K. pneumoniae* AzoA. Solving both structures under comparable experimental conditions enabled direct comparison of their active-site architectures, ligand-binding modes, and oligomeric assemblies. The structure was refined to 1.7 Å^2^ (Final R_work_ and R_free_ values of 16.5% and 19 %, respectively; Table S4) and contain one molecule in the asymmetric unit. However, crystallographic symmetry analysis indicates that AzoR forms a dimer with a total buried surface area of 3200 Å^2^ (Figure S5A). The structure highlights that unlike in AzoA, the cofactor FMN is solvent exposed (Figure S5B).

Attempts to soak these AzoA crystals, in mother liquor containing either NADPH or menadione lead to rapid dissolution of the crystals. We subsequently determined the structure of *K. pneumoniae* AzoA in the presence of Red 40 (PDB 7N2W; Table S4) using crystals grown via co-crystallization under conditions different to those of the holoenzyme. The structure was determined via molecular replacement using the holo-enzyme structure as the model and refined to a maximum resolution of 2.65 Å (Final R_work_ and R_free_ values of 24.9% and 30.2%, respectively; Table S4) with 4 protomers within the asymmetric unit. The assembly of the AzoA bound to Red 40 organizes a distinct dimeric interface compared to the holoenzyme and buries 3300 Å^2^ solvent exposed surface area (Figures 5A). The overall structure of each AzoA protomer remains consistent with that of the AzoA holoenzyme structure described previously [root-mean square deviation (RMSD) ~0.4 Å over 154 aligned Cα atoms]. The most significant change in this structure is that AzoA forms two distinct homodimers in the asymmetric unit (Figure 5A) rather than a dimer of dimers. DLS experiments with purified AzoA confirmed dissociation of AzoA tetramers into dimers in solution upon addition of either Red 40 or menadione (Figures 5B and S4). Inspection of the active site shows displacement of Gln145 (relocation of Cα >14 Å; Figure S7) and the L11 segment (aa 139-151) from the position seen in the holoenzyme structure (Figure 5C). In addition, in Red 40-bound AzoA, the 3_10_ helix within the L11 segment undergoes a conformational transition to a loop that closely resembles the corresponding conformations observed in *E. coli* AzoR (PDB 7N2X) and *S. typhimurium* AzoR (PDB 1T5B) (Figure S7). In the active site of the four protomers π-π stacking interactions are observed between the isoalloxazine rings of FMN and Red 40. In addition, Red 40 makes a network of polar interactions with surrounding residues (Figure 5D). The hydroxyl group of Red 40 forms hydrogen bonds with Arg60 and Tyr120 *in trans* (from the adjacent subunit). The two sulfonate groups of Red 40 contribute significantly to ligand recognition through electrostatic interactions within the active site. One sulfonate moiety is positioned near Arg183, forming a weak electrostatic contact in most protomers; however, in protomer C, a slight reorientation of the ligand enables formation of a direct salt bridge with the guanidinium group of Arg183. The second sulfonate group forms a salt bridge with His144 within the same protomer, providing an additional stabilizing interaction. The azo bond of Red 40 is positioned >4 Å from the N5 atom of FMN, exceeding the distance typically observed for direct electron transfer.^39, 40^ This geometry likely limits the efficiency of catalysis and is consistent with the poor turnover of Red 40 observed in our enzymatic assays.

### High-Affinity Binding of Red 40 to *K. pneumoniae* AzoA

To quantify the affinity of Red 40 for *K. pneumoniae* AzoA and characterize the thermodynamic basis of the interaction, we performed isothermal titration calorimetry (ITC). Successive injections of Red 40 into a solution of purified AzoA generated reproducible heat changes indicative of a direct binding interaction between the ligand and enzyme. The resulting thermodynamic binding isotherms were analyzed using the MicroCal PEAQ-ITC software package and were best described by a one-site binding model, consistent with a single predominant binding event under the experimental conditions employed.

Analysis of the ITC data revealed that Red 40 binds AzoA with high affinity, yielding an apparent dissociation constant (K_d_) of 27.1 ± 5.2 nM (Figure 6). This nanomolar affinity is consistent with the extensive network of protein-ligand interactions observed in the co-crystal structure, including hydrophobic packing, π-stacking interactions with the FMN cofactor, and multiple polar contacts within the active site. The exceptionally tight binding of Red 40 contrasts with its poor catalytic turnover, supporting the hypothesis that Red 40 behaves primarily as a high-affinity ligand, and potentially as an inhibitor, rather than as an efficiently reduced substrate. Together, the ITC and structural data demonstrate that AzoA recognizes Red 40 with remarkable affinity and provide insight into the molecular basis of this interaction.

**Figure 6.**
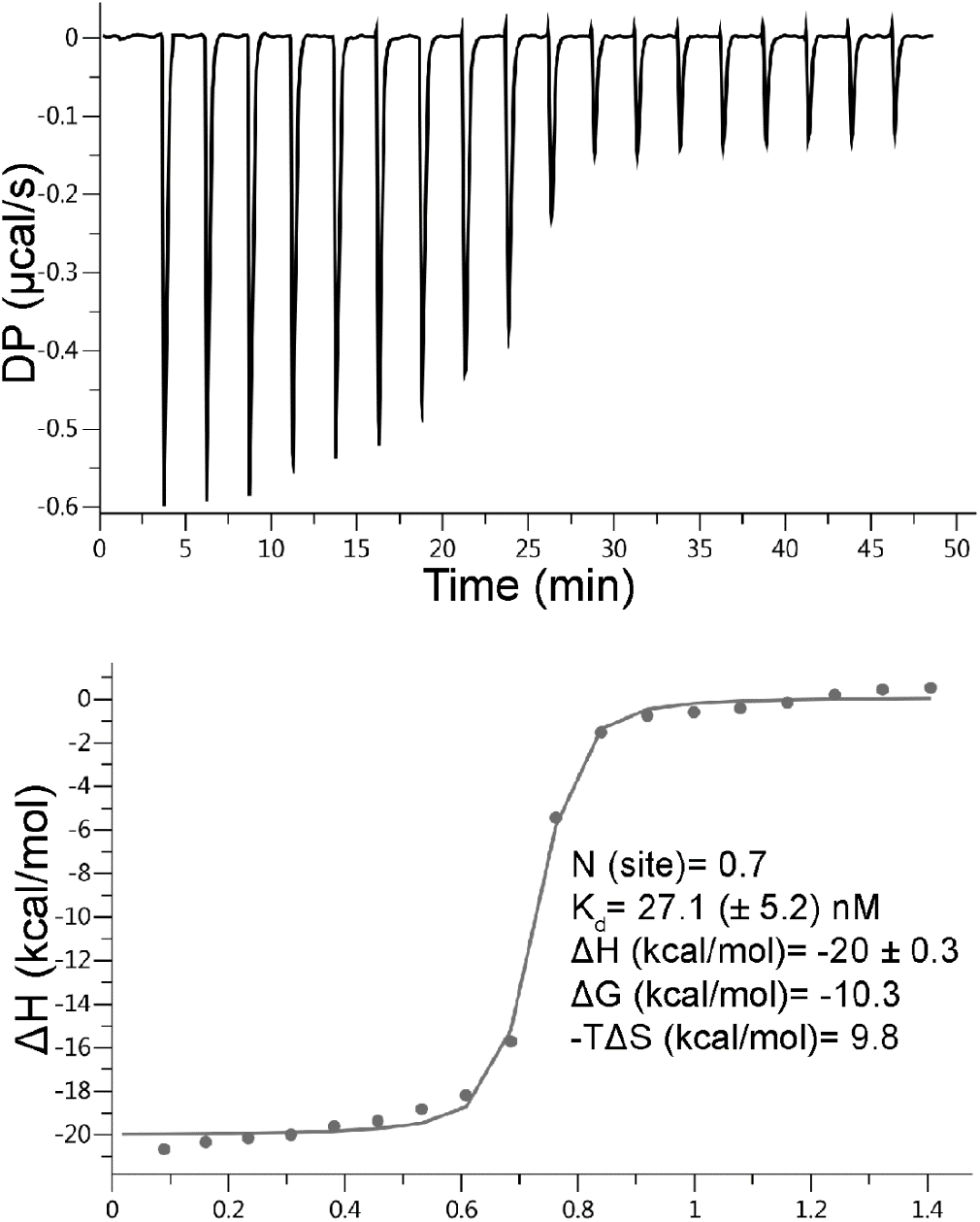
Binding of Red 40 to *Klebsiella pneumoniae* AzoA measured by isothermal titration calorimetry (ITC). (Upper panel) Representative ITC thermogram showing the raw heat signal generated by sequential injections of 150 μM Red 40 into a sample cell containing 10 μM AzoA (Experimental Methods). Each peak corresponds to the heat released or absorbed following a single ligand injection, and the integrated peak area reflects the heat associated with each binding event. **(Lower panel)** Binding isotherm obtained by integrating the heat from each injection and plotting the corrected heat change as a function of the Red 40 molar ratio. The data were fitted to a one-site binding model to determine the binding parameters. The fitted stoichiometry (N), dissociation constant (K_d_), enthalpy change (ΔH), Gibbs free energy change (ΔG), and entropic contribution (−TΔS) are indicated.

## DISCUSSION

### Diversity among azoreductases

Due to the remarkable diversity in protein sequence, phylogeny, and substrate profiles among azoreductases, identifying them via bioinformatic analysis proves challenging. To biochemically and structurally characterize representative family members, we analyzed the range of azoreductase sequences present within the gut microbiome.

Our analysis revealed that 76% of microbiome-associated azoreductase-like sequences fell into three main groups (clusters 1, 3, and 6; Figure 1A). Notably, structural information was available exclusively for proteins within Cluster 1. Of the 116 metagenomic sequences analyzed, only 49 exhibited high sequence similarity to a protein with an experimentally determined structure (E-value < 1 × 10^−20^), and all of these sequences belonged to Cluster 1 (Figure 1C). These publicly available structures are annotated as FMN-dependent NADH azoreductases from organisms such as *Bacillus* sp. or *Pseudomonas putida*. Notably, this type of azoreductase has recently been identified as a general reductase with a broad substrate profile, showing the highest activity with physiological quinone substrates.^18^

### Biochemical and Enzymatic Characterization of *K. pneumoniae* AzoA

*Klebsiella pneumoniae* is a Gram-negative bacterium, commonly found in the mucosal membranes of the oral and gastrointestinal tracts.^28^ Its colonization in the gut has been implicated as a causative agent of Inflammatory Bowel Disease (IBD) in mouse models.^41, 42^ Although *K. pneumoniae* strains were shown to reduce azo dyes over three decades ago,^43^ the specific enzymes responsible remain poorly understood. An azoreductase termed AzoK (52% identical to *K. pneumoniae* AzoA) has been partially characterized from *K. pneumoniae*,^44^ but its study was limited to methyl orange reduction, and its nicotinamide cofactor selectivity was not assessed. This highlights the gap in knowledge regarding azoreduction mechanisms in this organism.

Consistent with other Cluster 1 azoreductase-like enzymes,^19, 33^ the UV-Vis spectrum of purified recombinant AzoA (Figure 2B) displayed characteristic flavoprotein absorbance maxima at 360 and 450 nm,^45^ consistent with the presence of a bound flavin cofactor. Like the majority of bacterial azoreductases,^19, 24, 46^ AzoA incorporates non-covalently bound FMN, which is essential for electron transfer (Figure 4).^24^ Importantly, a key difference emerged regarding cofactor selectivity. Most Cluster 1 azoreductases exhibit a preference for either NADH or NADPH as electron donors.^33, 46^ In contrast, AzoA is reduced at comparable rates in the presence of either NADH or NADPH (Figure 3A).

### Substrate Profiling and Functional Hypothesis

When screened against a range of azo and quinone substrates, including common AFCs, prodrugs, and industrial dyes, AzoA showed relatively poor activity against all tested azo substrates. However, it reduced quinones, such as menadione, at a significantly faster rate (Figure 3B and Table S3). This substrate preference is consistent with data from other azoreductase-like enzymes within Cluster 1, specifically those from *Pseudomonas* species.^18, 47^ Similar to enzymes from *P. aeruginosa*, AzoA exhibited selectivity in quinone reduction, reducing benzoquinone approximately 100 times slower than the naphthoquinone menadione. This suggests that the primary physiological role of AzoA may be closer to that of a quinone oxidoreductase than a dedicated azoreductase.

### Contextualizing AzoA Activity with Gene Expression Data

A recently published study by Xie and colleagues^48^ provides further context for azoreduction in *K. pneumoniae*. This work demonstrated that AzoK (but not AzoA) was significantly upregulated during growth on the azo dye Reactive Black 5. Given that the expression of multiple azoreductases in a single organism is often dependent on the specific azo dye present in the growth medium,^43^ this finding does not definitively exclude AzoA from its role in azo reduction. However, in combination with the weak azo-reducing activity observed here, it strongly supports the hypothesis that azoreduction is not the primary function of AzoA.

The study by Xie and colleagues^48^ further highlighted the significant upregulation of: **1**) a *K. pneumoniae* homolog of WrbA, an NAD(P)H quinone oxidoreductase that shares the same short-flavodoxin-like fold as AzoA but does not reduce azo dyes;^34^ and **2**) membrane-bound NADH-quinone reductases, consistent with reports that homologs in *Shewanella* species can reduce Reactive Black 5.^49^ Together these transcriptional changes observed during azo dye exposure suggest that multiple, highly specific reductases, potentially including quinone oxidoreductases, contribute to the response of the bacterium.

### An Alternative Hypothesis for Azo Reduction

Recent studies have proposed an alternative, enzyme-independent pathway for azo compound reduction in the gut. ^50, 51^ Wolfson, <u>Hitchings,</u> and colleagues demonstrated that hydrogen sulfide (H_2_S), produced by the gut microbiome and present at millimolar concentrations (0.5-3 mM) in the intestinal lumen, can chemically reduce azo dyes.^50^ This chemical reduction mechanism is further supported by evidence from *E. coli*, where azoreductase metabolism is regulated by the Fumarate and Nitrate Reduction (FNR) regulator.^51^ The FNR regulator controls genes involved in H_2_S production from cysteine, suggesting a potential regulatory link between enzymatic and non-enzymatic pathways for azo compound reduction.

### Red 40-driven conformational change in AzoA

The overall structure of the AzoA holoenzyme is consistent with other Cluster 1 enzymes, such as the *E. coli* azoreductase (50% identical to PDB 7N2X and 2Z9B^52^ with a TM score^53^ of 0.89 across 199 residues). The primary difference observed in the AzoA structure is the reordering of the L11 segment (aa 139-151 in AzoA). In *E. coli* AzoR, this loop adopts a loop conformation (Figure S7); however, in the FMN bound AzoA structure, the amino acid segment forms a 3_10_ helix (Figures 4C and S7).

The formation of the 3_10_ helix in AzoA repositions the side chain of Gln145 close to the N10 atom of the FMN cofactor (Figure 4C). Crucially, this positioning precludes the binding of the nicotinamide group of NADH/NADPH in the required location for electron transfer (within 3.8 Å) of the N5 atom of FMN),^54^ and may suggest that the holo-enzyme is inactive in this state (Figures 4C).

Comparing AzoA with related azoreductases (e.g., *E. coli* AzoR; PDB 7N2X and *Salmonella typhimurium* AzoR; PDB 1T5B) shows the fold in the L11 segment is the only significant structural difference between the enzymes (Figure S7). Sequence alignments (Figure 7A) further suggest that substitutions at Gly146 and Thr148 in AzoA, corresponding to Glu146 and Pro148 in the related enzymes, may contribute to the unique conformation of the AzoA loop.^57^

**Figure 7.**
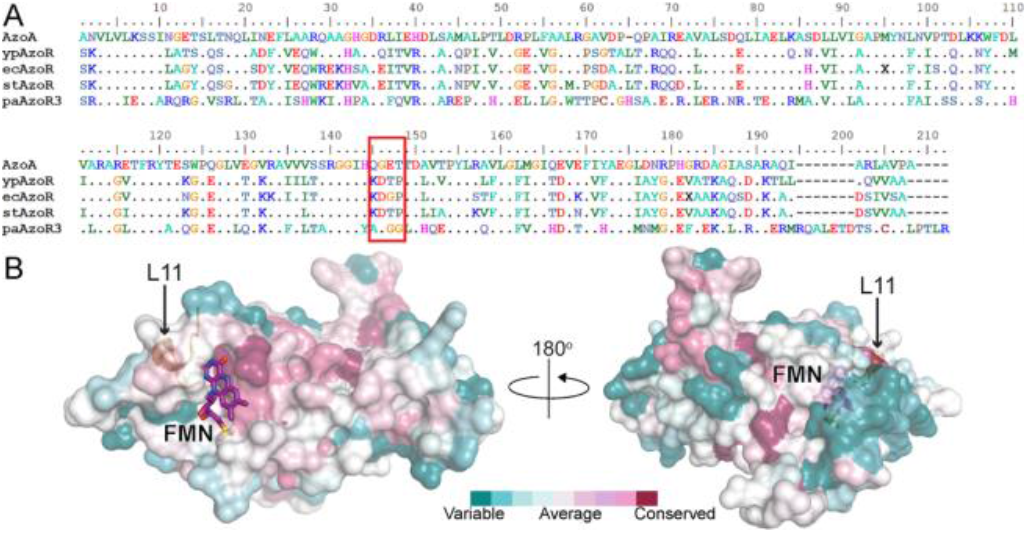
Sequence conservation of bacterial azoreductases. (A) Sequence alignment of K. pneumoniae AzoA and structurally characterized azoreductases. The L11 segment, which forms a 3_10_ helix in *K. pneumoniae* AzoA holoenzyme (Figures 4C and S7) and known for its sequence variability, is outlined in red. The alignment was performed using MUSCLE.^55^ (B) Surface representation of *K. pneumoniae* AzoA showing primary sequence conservation. Primary sequence conservation was mapped on the surface using ConSurf server.^56^ The color-coded bar indicates the level of conservation, ranging from variable (turquoise) to highly conserved (purple). Part of the L11 segment is colored gold to show its position and secondary structure within the AzoA holoenzyme. The FMN cofactor in the active site is shown as in Figure 4B.

Another key difference in the quaternary structure of AzoA is that the AzoA holoenzyme crystallizes as a homotetramer (a dimer of dimers; Figures 4A and 4B), whereas the *E. coli* (Figure S5) and *S. typhimurium* enzymes form strictly homodimeric structures. Tetramerization engages the L11 segment, which forms a 3_10_ helix in AzoA and contributes to the interface between the two dimers. This arrangement prevents the L11 segment from adopting the active conformation seen in the *E. coli* and *S. typhimurium* structures without disrupting the dimer-dimer interface. While some other Cluster 1 enzymes, like *P. aeruginosa* AzoR1 (38% identical to AzoA), form dimers of dimers in solution,^21^ their analogous L11 segment is not part of the interface, and therefore their activity is not constrained by the L11 conformation.

Unlike other Cluster 1 enzymes, which maintain a rigid active site upon ligand binding,^47, 52^ the dissolution of AzoA holo-enzyme crystals when a nicotinamide cofactor or electron acceptor is added strongly suggests that substrate binding induces a significant structural rearrangement. This supports the model where substrate binding displaces the residues forming the 3_10_ helix from the active site and disrupts the dimer-dimer interface of the tetramer (Figure 4B). Furthermore, crystals soaked in NADPH lose their yellow color, confirming FMN reduction.

Comparison of the Red 40 bound AzoA structure with the related enzymes shows that the 3_10_ helix has been displaced by the Red 40 molecule, and the L11 segment now adopts a conformation consistent with the active loop structures in the *E. coli* and *S. typhimurium* enzymes (Figure S7).

### Activation Mechanism of AzoA and Analogy with SsuE

Although the FMN position remains largely unchanged (Figure 5C), the rearrangement of the L11 segment causes a significant repositioning of the His144 side chain (Cα displacement of 11 Å; Figure 5C). This interaction is critical because it is postulated to stabilize the negative charge localized on the N1–C2=O moiety of the FMN that occurs following hydride transfer from NAD(P)H to the N5 atom.^54^

Substrate-driven changes in oligomeric state have also been observed in the sulfur starvation utilization protein E (SsuE),^58, 59^ which delivers reduced FMN to SsuD. SsuE, a short-flavodoxin fold containing protein, as a holoenzyme forms a tetramer (dimer of dimers). Upon binding of its substrate (FMN), the SsuE tetramer dissociates into homodimers.^58^ Unlike AzoA, this shift in oligomerization state in SsuE does not involve large structural changes but instead serves to expose the surface required for SsuD interaction.^60^

### Red 40 binding to AzoA

The crystal structure confirms that the binding of the food colorant Red 40 to AzoA is structurally consistent with the binding of other azo dyes in cluster 1 azoreductases, such as balsalazide in AzoR1 from *P. aeruginosa* (PDB 3LT5),^24^ and Acid Red 88 in AzrC from *Bacillus* sp. B29 (PDB 3W7A).^61^ These structures indicate that the dye participates in π-π stacking interactions with the FMN cofactor and forms both hydrogen-bonding and van der Waals interactions with a conserved tyrosine residue (Tyr120 in AzoA, Tyr131 in AzoR1, and Tyr127 in AzrC), which has been implicated in determining substrate specificity.^25^

It is noteworthy that unlike the predominantly hydrophobic contacts seen with substrates like balsalazide and methyl red in *P. aeruginosa* AzoR1,^19, 24^ Red 40 forms a large number of polar interactions with AzoA (Figure 5D). The Red 40 binding mode most closely resembles the interaction of the sulfonated azo dye, Acid Red 88, with AzrC (PDB 3W7A).^61^ However, the polar contacts differ significantly. Acid Red 88 forms a single hydrogen bond with Asn104 in AzrC. In contrast, Red 40 adopts a distinct binding pose in protomer C, resulting in a more extensive interaction network within the active site (Figure 5D). This conformation supports the formation of two additional hydrogen bonds and two salt bridges, likely due to the orientation of the methoxy substituent, which enables enhanced contacts with neighboring residues.

The observed π-π stacking interactions with the isoalloxazine rings of FMN, together with extensive polar and electrostatic interactions involving AzoA side chains, suggest a strong binding affinity for Red 40. This observation was confirmed by Isothermal Titration Calorimetry (ITC) measurements (Figure 6), which showed Red 40 binds AzoA tightly with an affinity of 27.1 (± 5.2) nM. This strong binding raises the possibility that Red 40 may inhibit AzoA activity by interfering with binding of the nicotinamide cofactor; however, direct inhibition was not experimentally assessed in the present study.

While the azo bond is spatially close (~4 Å) to the N5 atom of FMN (Figure 5D), Red 40 does not bind in a planar conformation. This non-planar conformation may suggest that the hydrazo tautomer of Red 40 is the bound species, similar to balsalazide binding to AzoR1.^24^ If electron transfer proceeds via the carbonyl carbon of the naphthalene ring, as expected for the hydrazo tautomer, the process would be inefficient, as this functionality is too distant (>6 Å) from the N5 of FMN. This spatial limitation may be a contributing factor to the low rate of azoreduction catalyzed by AzoA.

### Conclusion

To contextualize AzoA within the broader gut microbiome azoreductase family, a sequence conservation map was overlaid onto the dimeric structure (Figure 7B). Given the known low overall sequence conservation in this enzyme family,^34^ it is expected that the overlaid map mostly shows low conservation. Notably, the residues surrounding the FMN binding pocket exhibited higher conservation, suggesting that the structural scaffold required for cofactor binding is retained while the rest of the sequence is variable. However, the sequence encompassing the L11 segment, which mediates substrate-driven activation, is not highly conserved.

Our findings support the view that azoreductases are a specialized subgroup within the broader NAD(P)H-dependent quinone oxidoreductase superfamily. Although these enzymes share a conserved overall fold, substantial sequence and structural diversity exists both within the azoreductase family and among related quinone oxidoreductases. The ligand-dependent oligomeric transition observed for *K. pneumoniae* AzoA suggests a potential regulatory mechanism linking substrate binding to enzyme activity. However, given the marked differences in oligomeric organization among related enzymes such as MdaB, ChrR, and WrbA,^62, 63^ it remains unclear whether this phenomenon is broadly conserved. Further structural and mechanistic studies will be required to determine the extent to which ligand-coupled oligomeric transitions contribute to regulation across this enzyme superfamily.

## EXPERIMENTAL METHODS

### Azoreductase Sequence Database

The search term “azoreductase” was entered into NCBI protein, and all results were downloaded. This collection of sequences was dereplicated at 95% identity using USEARCH8 cluster_fast^64^ at 95% identity. This database is referred heretofore as azoDB.

### Identification of Azoreductases in the Human Gut Microbiome

Whole-community DNA gut metagenome assemblies from the NIH Integrative Human Microbiome Project (iHMP) IBDMDB (https://ibdmdb.org/)^31^ were downloaded and served as representative human gut metagenomes.^65^ From these nucleotide assemblies, we generated protein calls from the official assemblies using Prodigal 2.6.1 with the -p meta flag,^66^ and this database of all human gut metagenome proteins will be referred to as gutDB. at an e-value threshold of 1–10^−30^ using USEARCH8 with the options -mid 95 -accel 0.8. The resulting azoreductases identified from human gut metagenomes were dereplicated at an identity of 95%, using UCLUST8’s cluster fast algorithm. The resulting unique human-gut representative azoreductase sequences are presented as nodes in Figure 1.^67^

### Construction of the Sequence Similarity Network (SSN)

Before generation of the SSN, azoreductases identified by the text search “azoreductase” from the Protein Data Bank^68^ were added to the file of unique human-gut representative azoreductase sequences. This database of human-gut associated azoreductases were searched against themselves at an e-value threshold of 1*10^−20^ using USEARCH8 to generate a sequence similarity network. This network was visualized and analysed in Cytoscape 3.8.0^69^ in an undirected network using the default layout. Betweenness centrality was calculated using Cytoscape’s “analyze network” function. Nodes were colored according to iHMP metadata of disease status by referencing a sequence’s presence in a patient and that patient’s disease population according to IBDMDB metadata. Ternary plot of cluster composition was generated using R version 4.0.3 and the ‘Ternary’ package.^70^

### Dynamic Light Scattering (DLS)

DLS was conducted using a DynaPro Plate Reader (Wyatt Technology) at 20 °C. Samples of *K. pneumoniae* AzoA alone, its complexes with substrates and lysozyme were incubated at 20 °C for 15 min and centrifuged at 20,000 x g for 10 min to remove any suspended particulates before pipetting into a 384-well plate (Corning). The samples (25 µL/well; either AzoA alone at 0.5 mM concentration or with 1.5 mM substrates) in protein buffer were illuminated by 830 nm wavelength solid-state laser. The time scale of scattered light intensity fluctuations during each measurement was evaluated by autocorrelation, from which the translational diffusion coefficient (DT) was calculated. Molecular weights were estimated from the hydrodynamic radius (Rh), which was derived from DT using the Stokes-Einstein equation. The data was analyzed by Dynamics 7.8 software (Wyatt Technology).

### Enzyme Characterization

The absorbance spectrum of the enzyme was collected (BioTek Synergy HTX) to determine flavin binding compared to absorbance spectra of FMN. Cofactor selection was determined using NADH and NADPH. Substrates were manually chosen to represent biologically relevant quinones, azo-prodrugs, and azo-dyes. Aerobic enzyme stocks were buffer-exchanged using PD-10 desalting columns (GE Healthcare). Anaerobic experiments were performed in a benchtop anaerobic chamber sparged with ultrapure N_2_ (99.9%, Airgas) flowed over a heated copper catalyst to remove oxygen impurities. Anaerobic buffer was prepared by boiling double-distilled water to two-thirds initial volume and immediately maintained under positive nitrogen pressure while adding buffer components and stored in sparged anaerobic culture bottles stoppered under positive N_2_ pressure. Anaerobic protein buffer was made at concentrations of 100 mM Tris-HCl pH 7.4 and 100 mM NaCl. Enzyme assays were performed in a benchtop anaerobic chamber (Atmosbag, Milipore Sigma) sparged three times with ultrapure N_2_ flowed over a heated copper catalyst. After addition of all assay components to clear 96-well untreated half area plates (Costar, Corning Inc.), plates were sealed with clear PCR film (Santa Cruz Biotechnology) and removed from anaerobic chamber for spectrophotometric measurement on a plate reader (BioTek Synergy HTX). Final concentrations of the activity assay were: 500 µM NADH, 50 µM FMN, 100 µM substrate, 100 mM Tris pH 7.4, 100 mM NaCl, and 5 µg/mL (~0.2 µM) protein. Enzyme activity was determined by rate of NADH loss at 340 nm measured in continuous mode and quantified by external standardization on a linear scale using Gen5 software (version 1.11.5).

## ASSOCIATED CONTENT

Structure factors and coordinates of *Klebsiella pneumoniae* FMN-dependent NADH-azoreductase, AzoA (UniProt accession ID A6TCS9) are deposited in the RCSB Protein Data Bank with accession codes 6DXP (FMN) and 7N2W (FMN and Red 40). Structure factors and coordinates of *E. coli* FMN-dependent NADH-azoreductase, AzoR (UniProt accession ID P41407) are deposited in the RCSB Protein Data Bank with the accession code 7N2X (FMN)

### Supporting Information

The supporting information describes phylogeny of previously characterized azoreductases (Figure S1 and table S1), experimental methods for cloning, purification (Figure S2), Isothermal Titration Calorimetry, list of expressed AzoA orthologs (Table S2), summary of activity screen (Table S3), co-crystallization conditions for three AzoA complexes, RCSB accession codes, methods for crystallographic data processing, refinements (summarized in Table S4), 6 additional figures (Figures S2-S7) list of unique sequences of azoreductases from the NCBI protein database (95% identity; Supplemental data 1) and amino acid sequences of 63 diverse azoreductases from the human gut microbiome (Supplemental data 2). This material is available free of charge via the Internet at http://pubs.acs.org.AUTHOR INFORMATION

## AUTHORS INFORMATION

### Notes

The authors declare no competing financial interest.

## ACKNOWLEDGMENTS

We thank Elena Fedorov and Dr. Daniel J. Nilson for co-crystallization assistance. This research used resources of the U.S. Department of Energy (DOE) Office of Science supported User Facilities at the Argonne National Laboratory (contract no. DE-AC02-06CH11357). We acknowledge the support of the staff members at the LRL-CAT beamline at Sector 31 of the Advanced Photon Source.

## ABBREVIATIONS

AFC: artificial food coloring
DLS: dynamic Light Scattering
DT: diffusion coefficient
IBD: inflammatory bowel disease
IMAC: immobilized metal affinity chromatography
ITC: isothermal titration calorimetry
PDB: protein Databank Rh hydrodynamic radius
RMSD: root mean square deviation
SEC: size exclusion chromatography
SSN: sequence similarity network.

## REFERENCES

(1) Gürses, A.; Açıkyıldız, M.; Güneş, K.; Gürses, M. S. Classification of dye and pigments. In Dyes and Pigments, Springer International Publishing, 2016; pp 31–45.

(2) Batada, A.; Jacobson, M. F. Prevalence of artificial food colors in grocery store products marketed to children. Clin Pediatr (Phila) 2016, 55 (12), 1113–1119.

(3) Barciela, P.; Perez-Vazquez, A.; Prieto, M. A. Azo dyes in the food industry: Features, classification, toxicity, alternatives, and regulation. Food and Chemical Toxicology 2023, 178, 113935.

(4) Lehmkuhler, A.; Miller, M. D.; Bradman, A.; Castorina, R.; Chen, M.-A.; Xie, T.; Mitchell, A. E. Levels of FD&C certified food dyes in foods commonly consumed by children. J Food Composit Anal 2022, 112, DOI: 10.1016/j.jfca.2022.104649.

(5) Stevens, L. J.; Burgess, J. R.; Stochelski, M. A.; Kuczek, T. Amounts of artificial food dyes and added sugars in foods and sweets commonly consumed by children. Clin Pediatr (Phila) 2015, 54 (4), 309–321.

(6) Wainwright, M.; Kristiansen, J. E. On the 75th anniversary of Prontosil. Dyes and Pigments 2011, 88 (3), 231–234.

(7) Khanna, R.; Marshall, J. K. Sulfasalazine and 5-Aminosalicylates for Ulcerative Colitis. In Crohn’s Disease and Ulcerative Colitis, Baumgart, D. C. Ed.; Springer International Publishing, 2017; pp 389–397.

(8) Shi, C.-W.; Asch, S. M.; Fielder, E.; Gelberg, L.; Brook, R. H.; Leake, B.; Shapiro, M. F.; Dowling, P.; Nichol, M. Usage patterns of over-the-counter phenazopyridine (pyridium). J Gen Intern Med 2003, 18 (4), 281–287. PMCID: 12709095.

(9) Colebrook, L.; Buttle, G. A. H.; O’Meara, R. A. Q. The mode of action of p-aminobenzenesulphonamide and prontosil in haemolytic Streptococcal infections. Lancet 1936, 2 (5910), 1323–1326.

(10) Crettaz, S.; Kämpfer, P.; Brüschweiler, B. J.; Nussbaumer, S.; Deflorin, O. Survey on hazardous non-regulated aromatic amines as cleavage products of azo dyes found in clothing textiles on the Swiss market. J Consum Prot Food Saf 2020, 15 (1), 49–61.

(11) Jones, R.; Ryan, A. J.; Wright, S. E. The metabolism and excretion of tartrazine in the rat, rabbit and man. Food Cosmet Toxicol 1964, 2, 447–452.

(12) Roxon, J. J.; Ryan, A. J. Reduction of water-soluble azo dyes by intestinal bacteria. Food Cosmet Toxicol 1967, 5, 367–369.

(13) Gingell, R.; Bridges, J. W.; Williams, R. T. Gut flora and the metabolism of prontosils in the rat. Biochem J 1969, 114 (1), 5P–6P.

(14) Goldin, B.; Gorbach, S. L. Alterations in fecal microflora enzymes related to diet, age, lactobacillus supplements, and dimethylhydrazine. Cancer 1977, 40 (S5), 2421–2426.

(15) Mutlu, H.; Geiselhart, C. M.; Barner-Kowollik, C. Untapped potential for debonding on demand: the wonderful world of azo-compounds. Mater Horiz 2018, 5 (2), 162–183.

(16) Cerón-Carrasco, J. P.; Jacquemin, D. Using theory to extend the scope of azobenzene drugs in chemotherapy: Novel combinations for a specific delivery. ChemMedChem 2021, DOI: 10.1002/cmdc.202100046.

(17) Heinken, A.; Hertel, J.; Acharya, G.; Ravcheev, D. A.; Nyga, M.; Okpala, O. E.; Hogan, M.; Magnúsdóttir, S.; Martinelli, F.; Nap, B.; et al. Genome-scale metabolic reconstruction of 7,302 human microorganisms for personalized medicine. Nat Biotechnol 2023, DOI: 10.1038/s41587-41022-01628-41580.

(18) Ryan, A.; Kaplan, E.; Nebel, J.-C.; Polycarpou, E.; Crescente, V.; Lowe, E.; Preston, G. M.; Sim, E. Identification of NAD(P)H quinone oxidoreductase activity in azoreductases from P. aeruginosa: azoreductases and NAD(P)H quinone oxidoreductases belong to the same FMN-dependent superfamily of enzymes. PLoS ONE 2014, 9 (6), e98551.

(19) Romero, E.; Savino, S.; Fraaije, M. W.; Loncar, N. Mechanistic and crystallographic studies of azoreductase AzoA from Bacillus wakoensis A01. ACS Chem Biol 2020, 15 (2), 504–512.

(20) Ito, K.; Nakanishi, M.; Lee, W. C.; Sasaki, H.; Zenno, S.; Saigo, K.; Kitade, Y.; Tanokura, M. Three-dimensional structure of AzoR from Escherichia coli. An oxidereductase conserved in microorganisms. J Biol Chem 2006, 281 (29), 20567–20576. PMCID: 16684776.

(21) Wang, C. J.; Hagemeier, C.; Rahman, N.; Lowe, E.; Noble, M.; Coughtrie, M.; Sim, E.; Westwood, I. Molecular cloning, characterisation and ligand-bound structure of an azoreductase from Pseudomonas aeruginosa. J Mol Biol 2007, 373 (5), 1213–1228. PMCID: 17904577.

(22) Chen, H.; Wang, R. F.; Cerniglia, C. E. Molecular cloning, overexpression, purification, and characterization of an aerobic FMN-dependent azoreductase from Enterococcus faecalis. Protein Expr. Purif. 2004, 34 (2), 302–310. PMCID: 15003265.

(23) Binter, A.; Staunig, N.; Jelesarov, I.; Lohner, K.; Palfey, B. A.; Deller, S.; Gruber, K.; Macheroux, P. A single intersubunit salt bridge affects oligomerization and catalytic activity in a bacterial quinone reductase. FEBS J 2009, 276 (18), 5263–5274. PMCID: 19682074.

(24) Ryan, A.; Laurieri, N.; Westwood, I.; Wang, C. J.; Lowe, E.; Sim, E. A novel mechanism for azoreduction. J Mol Biol 2010, 400 (1), 24–37. PMCID: 20417637.

(25) Wang, C. J.; Laurieri, N.; Abuhammad, A.; Lowe, E.; Westwood, I.; Ryan, A.; Sim, E. Role of Tyrosine 131 in the active site of paAzoR1, an azoreductase with specificity for the inflammatory bowel disease pro-drug balsalazide. Acta Cryst 2010, F66 (1), 2–7.

(26) Ryan, A. Azoreductases in drug metabolism. Brit J Pharm 2017, 174 (14), 2161–2173.

(27) Chalansonnet, V.; Mercier, C.; Orenga, S.; Gilbert, C. Identification of Enterococcus faecalis enzymes with azoreductases and/or nitroreductase activity. BMC Microbiol 2017, 17 (1), 126. PMCID: 28545445PMC5445473.

(28) Gorrie, C. L.; Mirčeta, M.; Wick, R. R.; Edwards, D. J.; Thomson, N. R.; Strugnell, R. A.; Pratt, N. F.; Garlick, J. S.; Watson, K. M.; Pilcher, D. V.; et al. Gastrointestinal carriage is a major reservoir of Klebsiella pneumoniae infection in intensive care patients. Clin Infect Dis 2017, 65 (2), 208–215.

(29) Martin, R. M.; Cao, J.; Brisse, S.; Passet, V.; Wu, W.; Zhao, L.; Malani, P. N.; Rao, K.; Bachman, M. A. Molecular epidemiology of colonizing and infecting isolates of Klebsiella pneumoniae. mSphere 2016, 1 (5), e00261–00216.

(30) Sayers, E. W.; Bolton, E. E.; Brister, J. R.; Canese, K.; Chan, J.; Comeau, D. C.; Connor, R.; Funk, K.; Kelly, C.; Kim, S.; et al. Database resources of the national center for biotechnology information. Nucleic Acids Res 2022, 50 (D1), D20–d26. PMCID: 34850941PMC8728269.

(31) Lloyd-Price, J.; Arze, C.; Ananthakrishnan, A. N.; Schirmer, M.; Avila-Pacheco, J.; Poon, T. W.; Andrews, E.; Ajami, N. J.; Bonham, K. S.; Brislawn, C. J.; et al. Multi-omics of the gut microbial ecosystem in inflammatory bowel diseases. Nature 2019, 569 (7758), 655–662.

(32) Nakanishi, M.; Yatome, C.; Ishida, N.; Kitade, Y. Putative ACP phosphodiesterase gene (acpD) encodes an azoreductase. J Biol Chem 2001, 276 (49), 46394–46399. PMCID: 11583992.

(33) Ryan, A.; Wang, C. J.; Laurieri, N.; Westwood, I.; Sim, E. Reaction mechanism of azoreductases suggests convergent evolution with quinone oxidoreductases. Protein Cell 2010, 1 (8), 780–790. PMCID: 21203919.

(34) Crescente, V.; Holland, S. M.; Kashyap, S.; Polycarpou, E.; Sim, E.; Ryan, A. Identification of novel members of the bacterial azoreductase family in Pseudomonas aeruginosa. Biochem J 2016, 473 (5), 549–558.

(35) Ye, J.; Yang, H. C.; Rosen, B. P.; Bhattacharjee, H. Crystal structure of the flavoprotein ArsH from Sinorhizobium meliloti. FEBS Lett 2007, 581 (21), 3996–4000. PMCID: 17673204.

(36) Fekih, I. B.; Zhang, C.; Li, Y. P.; Zhao, Y.; Alwathnani, H. A.; Saquib, Q.; Rensing, C.; Cervantes, C. Distribution of Arsenic resistance genes in prokaryotes. Front Microbiol 2018, 9, DOI: 10.3389/fmicb.2018.02473, Review.

(37) Paez-Espino, A.; Nikel, P. I.; Chavarria, M.; de Lorenzo, V. ArsH protects Pseudomonas putida from oxidative damage caused by exposure to arsenic. Environ Microbiol 2020, 22 (6), 2230–2242.

(38) Krissinel, E.; Henrick, K. Inference of macromolecular assemblies from crystalline state. J Mol Biol 2007, 372 (3), 774–797. PMCID: 17681537.

(39) Faig, M.; Bianchet, M. A.; Talalay, P.; Chen, S.; Winski, S.; Ross, D.; Amzel, L. M. Structures of recombinant human and mouse NAD(P)H:quinone oxidoreductases: species comparison and structural changes with substrate binding and release. Proc Natl Acad Sci U S A 2000, 97 (7), 3177–3182. PMCID: 10706635PMC16212.

(40) Degtjarik, O.; Brynda, J.; Ettrichova, O.; Kuty, M.; Sinha, D.; Kuta Smatanova, I.; Carey, J.; Ettrich, R.; Reha, D. Quantum Calculations Indicate Effective Electron Transfer between FMN and Benzoquinone in a New Crystal Structure of Escherichia coli WrbA. J Phys Chem B 2016, 120 (22), 4867–4877. PMCID: 27183467.

(41) Zhang, Q.; Su, X.; Zhang, C.; Chen, W.; Wang, Y.; Yang, X.; Liu, D.; Zhang, Y.; Yang, R. Klebsiella pneumoniae induces inflammatory bowel disease through Caspase-11–Mediated IL18 in the gut epithelial cells. Cell Mol Gastroenterol Hepatol 2023, 15 (3), 613–632.

(42) Federici, S.; Kredo-Russo, S.; Valdés-Mas, R.; Kviatcovsky, D.; Weinstock, E.; Matiuhin, Y.; Silberberg, Y.; Atarashi, K.; Furuichi, M.; Oka, A.; et al. Targeted suppression of human IBD-associated gut microbiota commensals by phage consortia for treatment of intestinal inflammation. Cell 2022, 185 (16), 2879–2898.e2824. PMCID: 35931020.

(43) Brown, J. P. Reduction of polymeric azo and nitro dyes by intestinal bacteria. App Env Micro 1981, 41 (5), 1283–1286.

(44) Dixit, S.; Garg, S. Enzymatic degradation of sulphonated azo dye using purified azoreductase from facultative Klebsiella pneumoniae. Folia Microbiol 2021, 66, 79–85.

(45) Macheroux, P. UV-Visible spectroscopy as a tool to study flavoproteins. In Flavoprotein protocols, Methods Mol Biol, Vol. 131; Humana Press, 1999; pp 1–8.

(46) Chen, H.; Hopper, S. L.; Cerniglia, C. E. Biochemical and molecular characterization of an azoreductase from Staphylococcus aureus, a tetrameric NADPH-dependent flavoprotein. Microbiology 2005, 151 (5), 1433–1441. PMCID: 15870453.

(47) Gonçalves, A. M. D.; Mendes, S.; de Sanctis, D.; Martins, L. O.; Bento, I. The crystal structure of Pseudomonas putida AzoR: the active site revisited. FEBS J 2013, 280 (24), 6643–6657.

(48) Xie, X.; Zheng, H.; Zhang, Q.; Fan, J.; Liu, N.; Song, X. Cometabolic biodegradation of structurally discrepant dyestuffs by Klebsiella sp. KL-1: A molecular mechanism with regards to the differential responsiveness. Chemosphere 2022, 303, DOI: 10.1016/j.chemosphere.2022.135028.

(49) Imran, M.; Negm, F.; Hussain, S.; Ashraf, M.; Ashraf, M.; Ahmad, Z.; Arshad, M.; Crowley, D. E. Characterization and purification of membrane-bound azoreductase from azo dye degrading Shewanella sp. Strain IFN4. Clean Soil, Air, Water 2016, 44 (11), 1523–1530.

(50) Wolfson, S. J.; Hitchings, R.; Peregrina, K.; Cohen, Z.; Khan, S.; Yilmaz, T.; Malena, M.; Goluch, E. D.; Augenlicht, L.; Kelly, L. Bacterial hydrogen sulfide drives cryptic redox chemistry in gut microbial communities. Nat Metab 2022, 4 (10), 1260–1270.

(51) Pieper, L. M.; Spanogiannopoulos, P.; Volk, R. F.; Miller, C. J.; Wright, A. T.; Turnbaugh, P. J. The global anaerobic metabolism regulator fnr is necessary for the degradation of food dyes and drugs by Escherichia coli. bioRxiv 2022, DOI: 2022.2010.2025.513779.

(52) Ito, K.; Nakanishi, M.; Lee, W. C.; Zhi, Y.; Sasaki, H.; Zenno, S.; Saigo, K.; Kitade, Y.; Tanokura, M. Expansion of substrate specificity and catalytic mechanism of azoreductase by X-ray crystallography and site-directed mutagenesis. J Biol Chem 2008, 283 (20), 13889–13896. PMCID: 18337254.

(53) Zhang, Y.; Skolnick, J. Scoring function for automated assessment of protein structure template quality. Proteins 2004, 57 (4), 702–710. PMCID: 15476259.

(54) Fraaije, M. W.; Mattevi, A. Flavoenzymes: diverse catalysts with recurrent features. Trends Biochem Sci 2000, 25 (3), 126–132. PMCID: 10694883.

(55) Edgar, R. C. MUSCLE: multiple sequence alignment with high accuracy and high throughput. Nucleic Acids Res 2004, 32 (5), 1792–1797. PMCID: 15034147390337.

(56) Yariv, B.; Yariv, E.; Kessel, A.; Masrati, G.; Chorin, A. B.; Martz, E.; Mayrose, I.; Pupko, T.; Ben-Tal, N. Using evolutionary data to make sense of macromolecules with a “face-lifted” ConSurf. Protein Sci 2023, 32 (3), e4582. PMCID: 36718848PMC9942591.

(57) Huang, F.; Nau, W. M. A conformational flexibility scale for amino acids in peptides. Angewandte Chemie-International Edition 2003, 42 (20), 2269–2272, Article. PMCID: WOS:000183334500010.

(58) Driggers, C. M.; Dayal, P. V.; Ellis, H. R.; Karplus, P. A. Crystal structure of Escherichia coli SsuE: Defining a general catalytic cycle for FMN reductases of the flavodoxin-like superfamily. Biochemistry 2014, 53 (21), 3509–3519.

(59) Aloh, C. H.; Zeczycki, T. N.; Ellis, H. R. Oligomeric Changes Regulate Flavin Transfer in Two-Component FMN Reductases Involved in Sulfur Metabolism. Biochemistry 2023, 62 (18), 2751–2762. PMCID: 37651343.

(60) Dayal, P. V.; Singh, H.; Busenlehner, L. S.; Ellis, H. R. Exposing the Alkanesulfonate monooxygenase protein–protein interaction sites. Biochemistry 2015, 54 (51), 7531–7538.

(61) Yu, J.; Ogata, D.; Gai, Z.; Taguchi, S.; Tanaka, I.; Ooi, T.; Yao, M. Structures of AzrA and of AzrC complexed with substrate or inhibitor: insight into substrate specificity and catalytic mechanism. Acta Cryst 2014, D70 (2), 553–564.

(62) Kishko, I.; Carey, J.; Reha, D.; Brynda, J.; Winkler, R.; Harish, B.; Guerra, R.; Ettrichova, O.; Kukacka, Z.; Sheryemyetyeva, O.; et al. 1.2 A resolution crystal structure of Escherichia coli WrbA holoprotein. Acta Crystallogr D Biol Crystallogr 2013, 69 (Pt 9), 1748–1757. PMCID: 23999298.

(63) Eswaramoorthy, S.; Poulain, S.; Hienerwadel, R.; Bremond, N.; Sylvester, M. D.; Zhang, Y. B.; Berthomieu, C.; Van Der Lelie, D.; Matin, A. Crystal structure of ChrR--a quinone reductase with the capacity to reduce chromate. PLoS One 2012, 7 (4), DOI: 10.1371/journal.pone.0036017. PMCID: 22558308PMC3338774.

(64) Edgar, R. C. Search and clustering orders of magnitude faster than BLAST. Bioinformatics 2010, 26 (19), 2460–2461.

(65) iHMP. The integrative human microbiome project: Dynamic analysis of microbiome-host omics profiles during periods of human health and disease. Cell Host & Microbe 2014, 16 (3), 276–289.

(66) Hyatt, D.; Chen, G.-L.; Locascio, P. F.; Land, M. L.; Larimer, F. W.; Hauser, L. J. Prodigal: prokaryotic gene recognition and translation initiation site identification. BMC Bioinformatics 2010, 11, DOI: 10.1186/1471-2105-1111-1119. PMCID: 20211023.

(67) Edgar, R. C. Search and clustering orders of magnitude faster than BLAST. Bioinformatics 2010, 26 (19), 2460–2461. PMCID: 20709691.

(68) Berman, H. M.; Westbrook, J.; Feng, Z.; Gilliland, G.; Bhat, T. N.; Weissig, H.; Shindyalov, I. N.; Bourne, P. E. The Protein Data Bank. Nucleic Acids Res 2000, 28 (1), 235–242.

(69) Shannon, P.; Markiel, A.; Ozier, O.; Baliga, N. S.; Wang, J. T.; Ramage, D.; Amin, N.; Schwikowski, B.; Ideker, T. Cytoscape: a software environment for integrated models of biomolecular interaction networks. Genome Res 2003, 13 (11), 2498–2504. PMCID: 14597658PMC403769.

(70) Smith, M. Ternary: An R Package for Creating Ternary Plots. Zenodo 2017, DOI: 10.5281/zenodo.1068996.

