## Supporting Information for "Ligand binding remodels the oligomeric state and active site of *Klebsiella pneumoniae* azoreductase"

### Experimental Methods

**Bayesian Phylogenetic Analysis** Enzymes previously investigated for azoreduction were identified from the literature and retrieved from RefSeq. Phylogenetic analysis was performed using the MrBayes algorithm implemented in Ugene 35.1 with the default parameters: poisson rate matrix and gamma rate variation model with 4 rate categories. Presence in a particular cluster was determined by mapping to the centroid sequence generated by clustering at 95% identity to produce NCBI_azoreductases_c95.fasta (supplemental data 1 and Table S1).

**Cloning and Protein Expression** Azoreductase sequences with the highest betweenness centrality were manually chosen to represent a diverse sequence space. These sequences were codon-optimized for *E. coli* expression and *de novo* synthesized at IDT DNA (Coralville, IA). Sequences were appended with an N-terminal $His_6$ tag and then inserted into the plasmid pSGC for expression using ligation-independent cloning.[1] Plasmids containing azoreductases were transformed into *E. coli* DH10b cultures for plasmid amplification then transformed into the *E. coli* BL21(DE3) pRIL expression strain and expressed in SeMet incorporation PASM-5052 media[2] grown in a large-scale expression (LEX) bioreactor system (Epiphyte3) maintained at 37° C.

**Protein Purification** Cells were harvested by centrifugation at 2000xg and suspended in buffer containing 10 mM HEPES (pH 7.6), 150 mM NaCl, 20 mM imidazole, and 1 mM β-mercaptoethanol (βME). Cells were lysed by sonication and insoluble cell precipitate was removed by centrifugation at 27,000xg. Soluble expressed proteins were purified by IMAC using a $His_{60}$ Ni Superflow resin (Takara) equilibrated with buffer containing 10 mM HEPES (pH 7.6), 150 mM NaCl, 1 mM βME and 20 mM imidazole. After lysate loading, the column was washed with the same buffer containing 200 mM imidazole to elute the protein and collected. The $His_6$ tag was removed by digestion overnight with TEV protease at 4° C. Preparatory SEC was performed with Superdex S200 PG resin (Cytiva) in buffer containing 10 mM HEPES pH 7.6 and

150 mM NaCl on an ÅKTA Xpress FPLC system and the fractions containing purified protein were collected and pooled. Purified enzymes were analyzed by SDS-PAGE for confirmation of overexpression of bands at the appropriate predicted molecular weight and purity. AzoA protein concentration was determined using the Bradford assay (Thermo Fisher).

**Isothermal Titration Calorimetry (ITC).** ITC experiments were performed using a MicroCal PEAQ-ITC instrument. AzoA and Red 40 were prepared in degassed buffer containing 20 mM HEPES (pH 7.6), 150 mM NaCl, and 1 mM β-mercaptoethanol (βME). The sample cell was loaded with 300 μL of 10 μM AzoA, and the injection syringe was filled with approximately 40 μL of 150 μM Red 40. The reference cell contained 300 μL of filtered (0.2 μm, Millipore) deionized water. Titrations were performed at 25 °C using 2 μL injections at 150 s intervals. The resulting binding isotherms were analyzed using one-site or two-site binding models with the MicroCal PEAQ-ITC Analysis software.

**Protein crystallization and crystal harvesting** Sitting drop vapor diffusion plates were set up at room temperature. Single crystals of *K. pneumoniae* AzoA holoenzyme were grown from a solution containing 368 μM protein in 10 mM HEPES, pH 7.6, and 150 mM NaCl with the following precipitant conditions 0.1 M MES pH 6.0, 25% w/v PEG 8000, and 0.2 M calcium acetate. Crystals of *K. pneumoniae* AzoA and Red 40 complex were obtained using solutions of 815 μM protein and 1.5 mM Red 40 in sitting drops against well solutions containing 1.26 M ammonium sulphate, 0.2 M NaCl and 0.1 M CHES pH 9.5 at 19 °C. Crystals of *K. pneumoniae* AzoA and Red 40 complex were cryoprotected by addition of 30% glycerol to the mother liquor prior to flash-cooling.

**Data collection and processing, structure determination, model building, refinement and analysis** Diffraction data for all crystals were collected with a CCD Rayonix MX-225 HE detector on beamline 31-ID at the Advanced Photon Source of Argonne National Laboratory. Data were integrated using MOSFLM[3] and scaled using SCALA.[4] Phasing of AzoA holoenzyme was performed via molecular replacement in PHASER[5] using the structure of *E. coli* azoreductase

(PDB 1V4B) [6] as the search model. Multiple rounds of refinement were then performed using REFMAC5[7] and PHENIX[8] interspersed with rounds of model building in COOT.[9] Processing and refinement statistics are provided in Table S4.

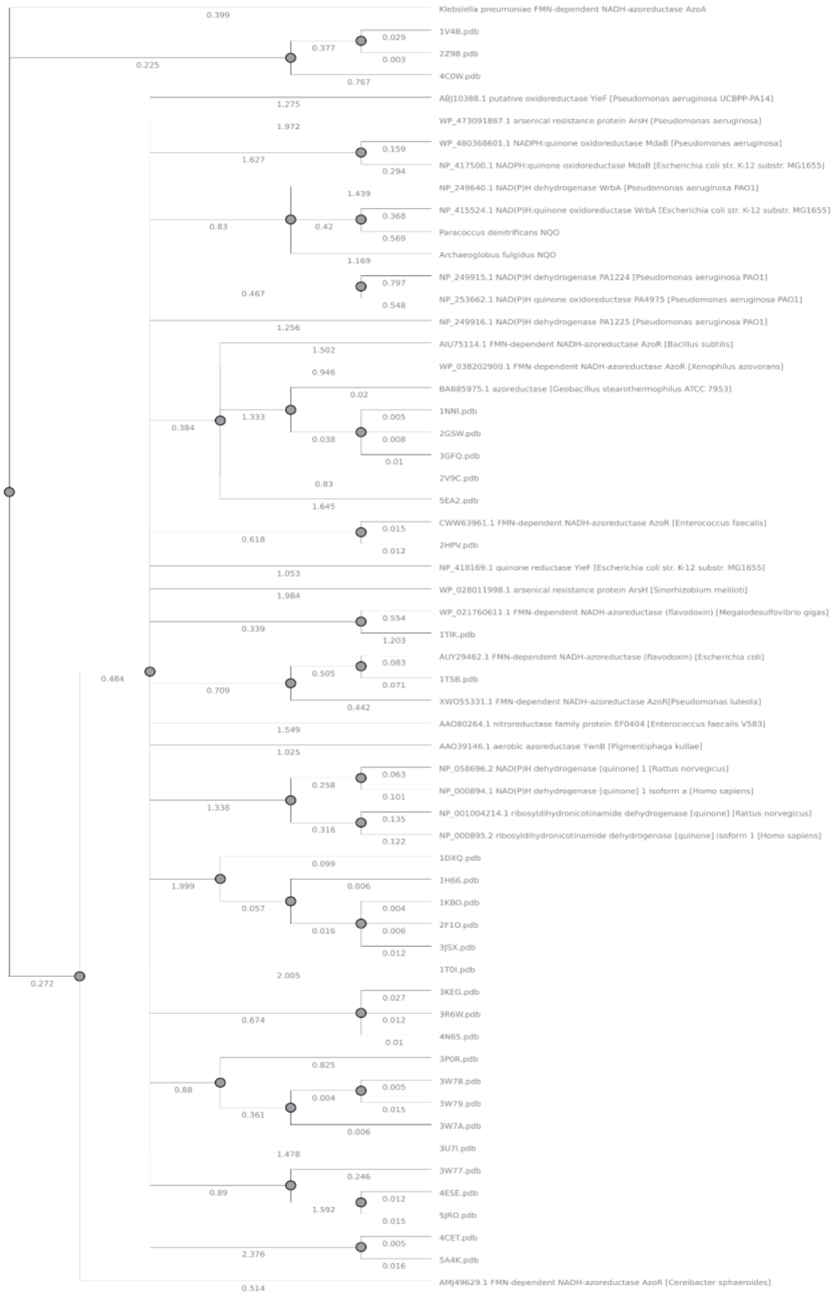

**Figure S1. Phylogeny of previously characterized azoreductases**. Bayesian phylogenetic analysis of previously characterized azoreductase-like enzymes. Sequences with experimentally determined structures deposited in the Protein Data Bank (PDB) are labeled with their PDB accession codes followed by the suffix ".pdb"; all other sequences are labeled using their RefSeq identifiers.

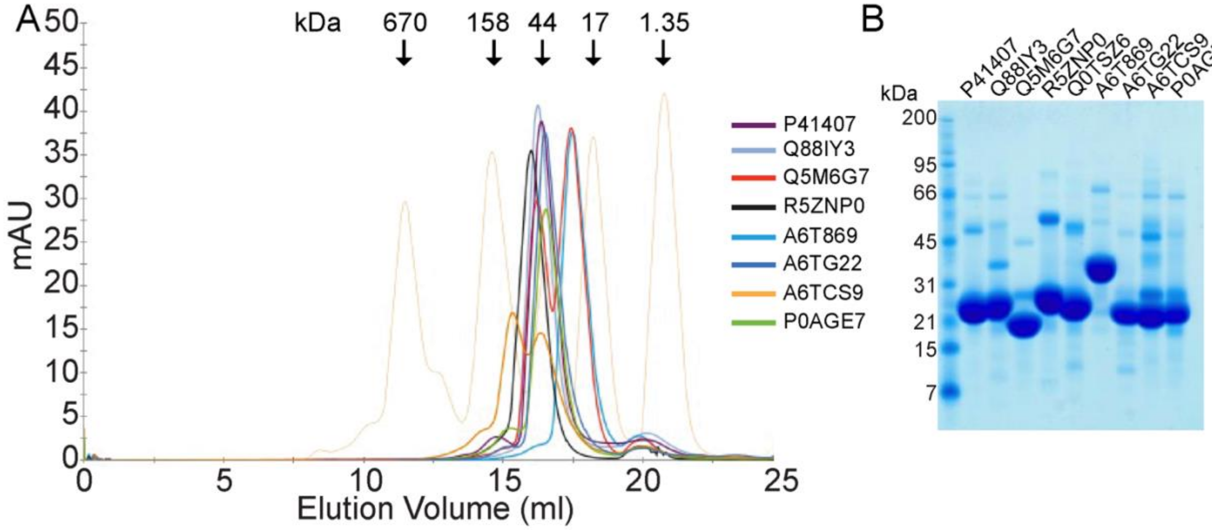

**Figure S2. Expression of AzoA orthologs.** (A) Size-exclusion chromatography (SEC) profiles of AzoA orthologs identified by their respective UniProt accession numbers. Protein elution was monitored by absorbance at 280 nm (mAU) using a calibrated Superdex 200 Increase 10/300 GL column under the conditions described in the Experimental Methods. The thin orange trace represents the elution positions of molecular weight standards, with the corresponding molecular masses indicated, obtained under identical buffer conditions. (B) Aliquots of nine orthologous AzoA were analyzed by SDS-PAGE. The positions and sizes (kDa) of marker proteins are indicated on the *left*.

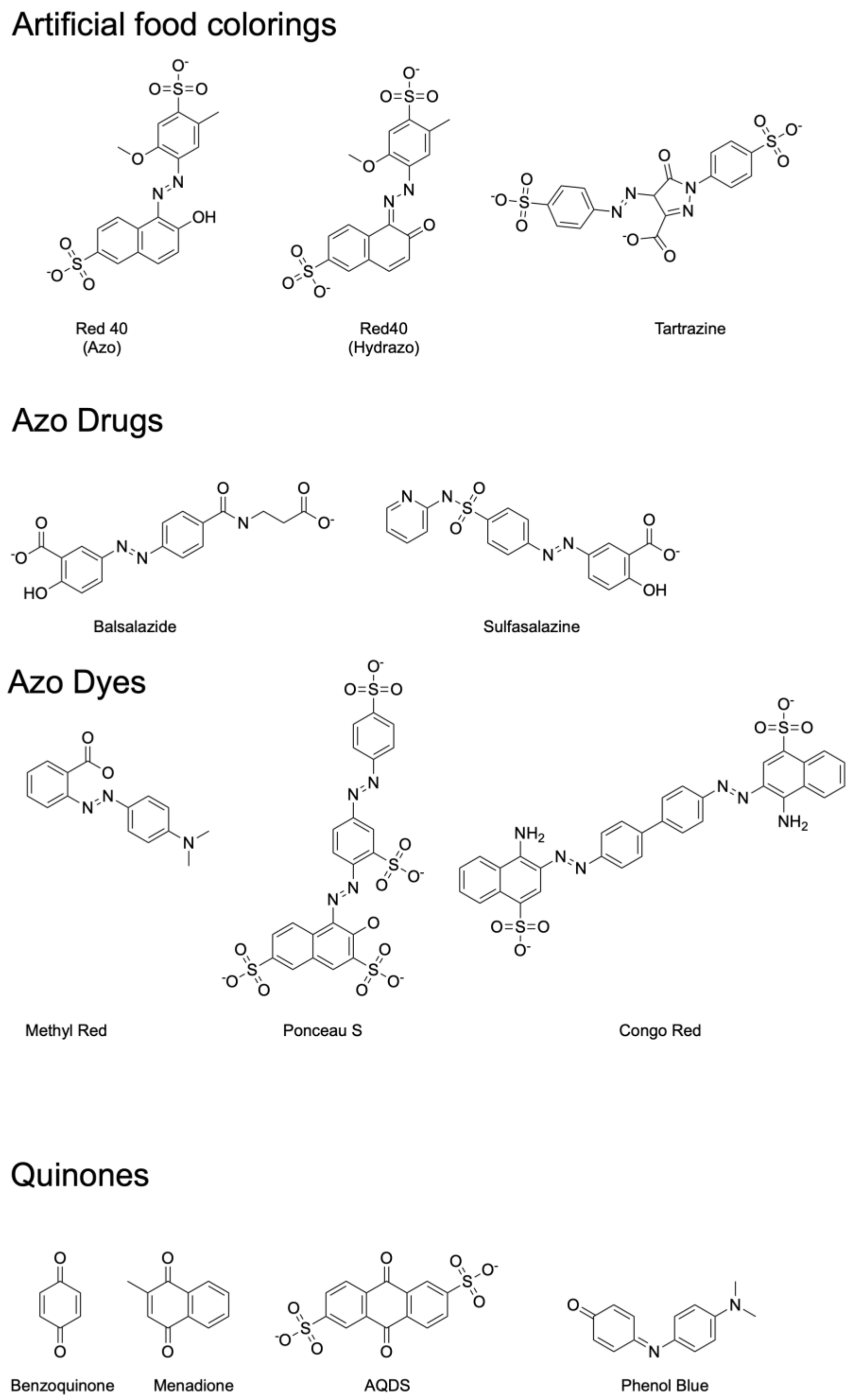

**Figure S3: AzoA substrates.** Schematic of the artificial food colorings (AFCs)/substrates (as indicated) used for characterization of AzoA.

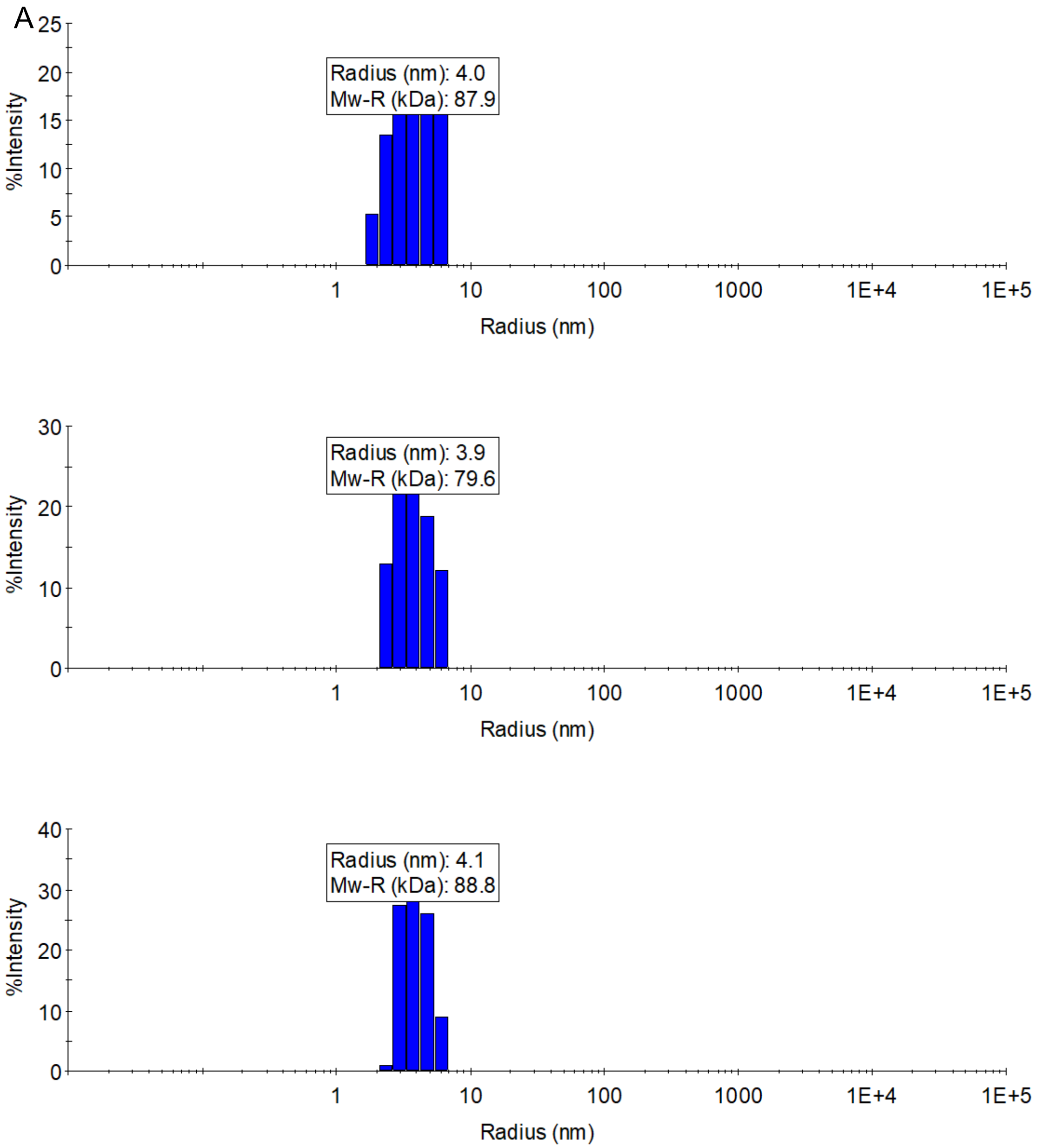
A
Radius (nm): 4.0
Mw-R (kDa): 87.9
%Intensity
Radius (nm)
Radius (nm): 3.9
Mw-R (kDa): 79.6
%Intensity
Radius (nm)
Radius (nm): 4.1
Mw-R (kDa): 88.8
%Intensity
Radius (nm)

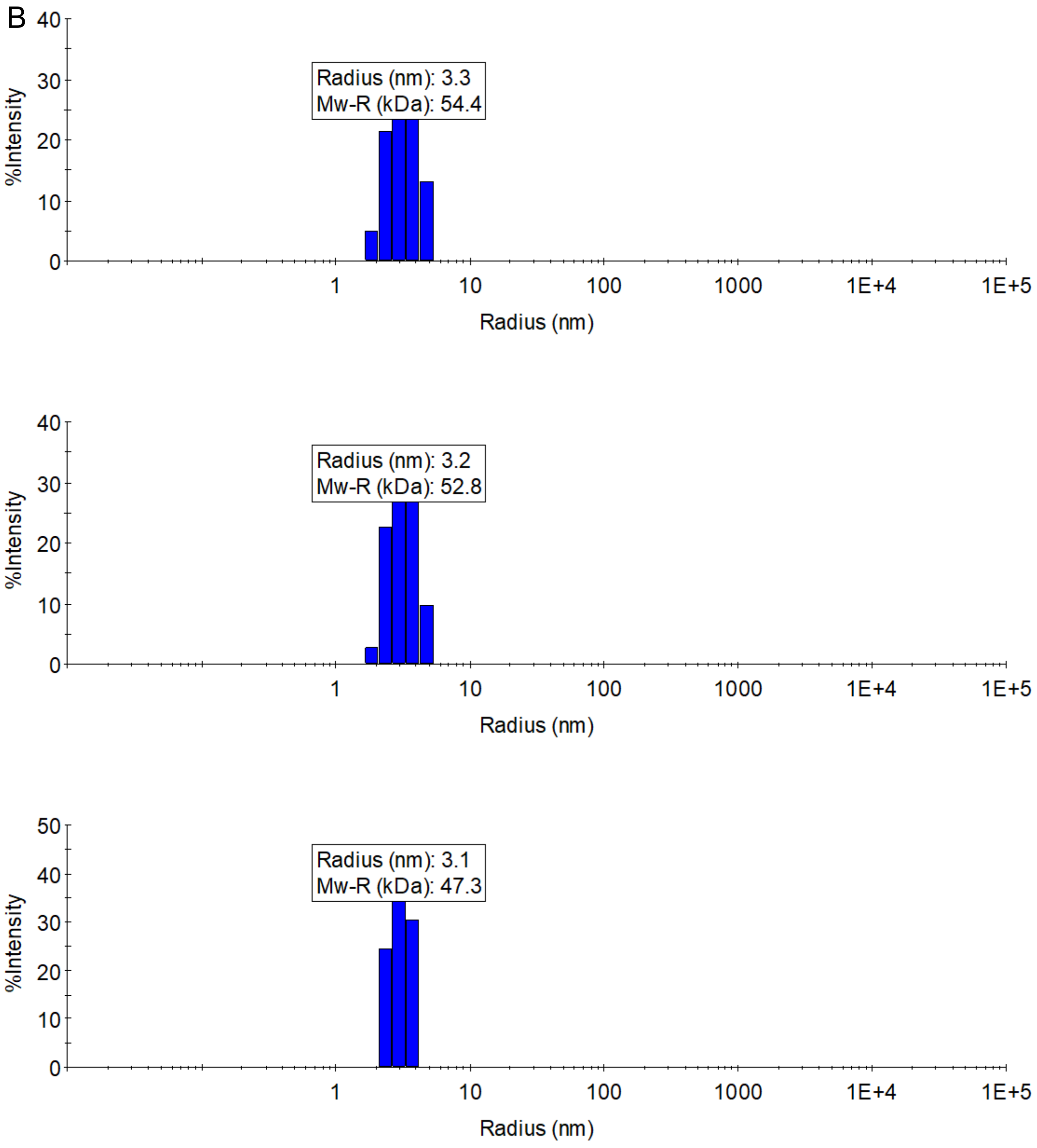
B
Radius (nm): 3.3
Mw-R (kDa): 54.4
%Intensity
Radius (nm)
Radius (nm): 3.2
Mw-R (kDa): 52.8
Radius (nm): 3.1
Mw-R (kDa): 47.3

D

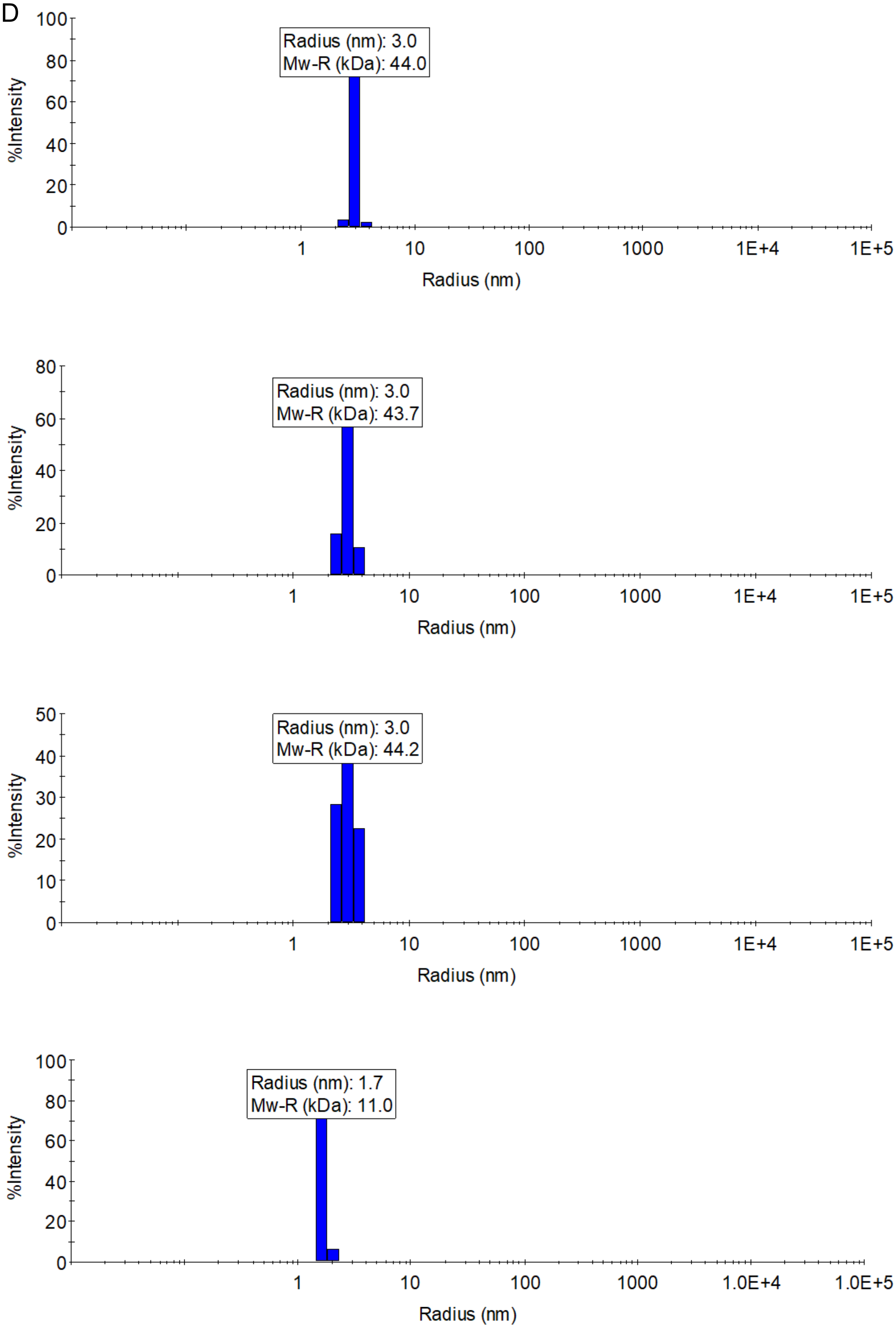
Radius (nm): 3.0
Mw-R (kDa): 44.0
%Intensity
Radius (nm)
Radius (nm): 3.0
Mw-R (kDa): 43.7
%Intensity
Radius (nm)
Radius (nm): 3.0
Mw-R (kDa): 44.2
%Intensity
Radius (nm)
Radius (nm): 1.7
Mw-R (kDa): 11.0
%Intensity
Radius (nm)

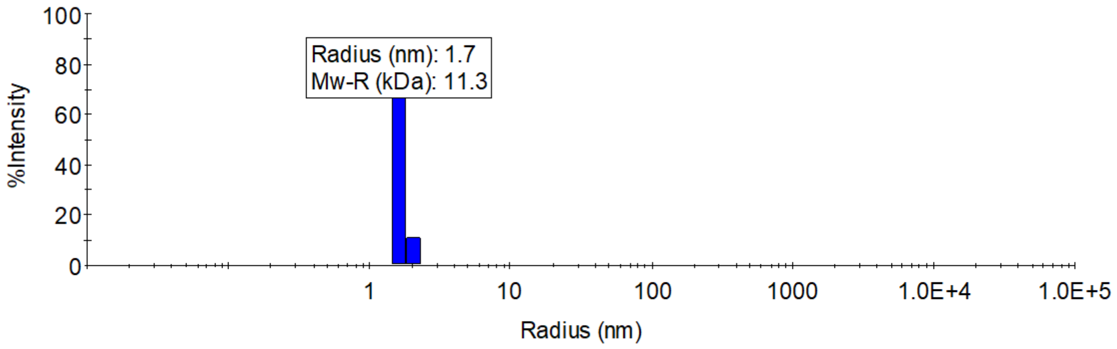

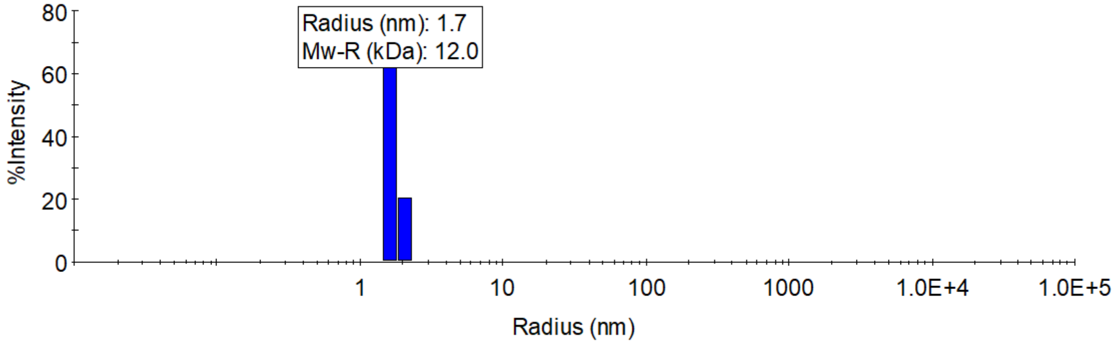

**Figure S4: Solution behavior of AzoA holoenzyme alone and in the presence of substrates.** Calculated apparent molecular weights (Figure 2D) of *K. pneumoniae* AzoA (A), AzoA in the presence of Red 40 (B) and Menadione (C) along with lysozyme (D) as a reference determined from hydrodynamic radii using Dynamic Light Scattering (Experimental Methods).

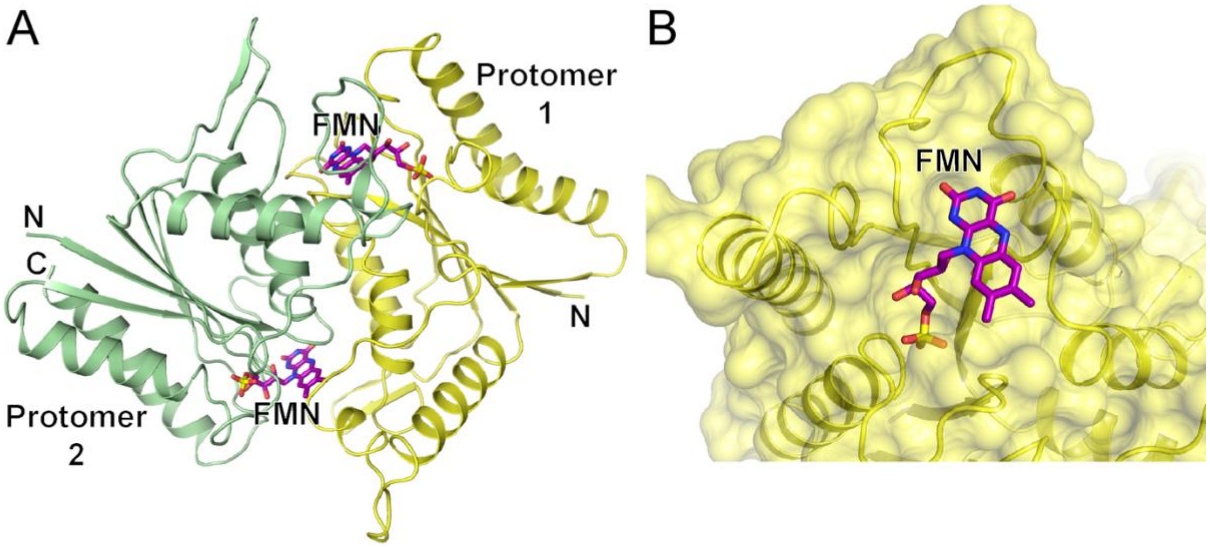

**Figure S5: The structure of *E. coli* AzoR. A)** AzoR dimer shown as in Figure 1A a ribbon with arrows for β-strands and wide ribbons for helices. Two symmetry related protomers of the dimer are color coded in yellow and green. Bound FMNs are shown in stick representations and colored purple. **B)** Close-up view of an AzoR active site showing bound flavin mononucleotide (FMN) cofactor is solvent exposed.

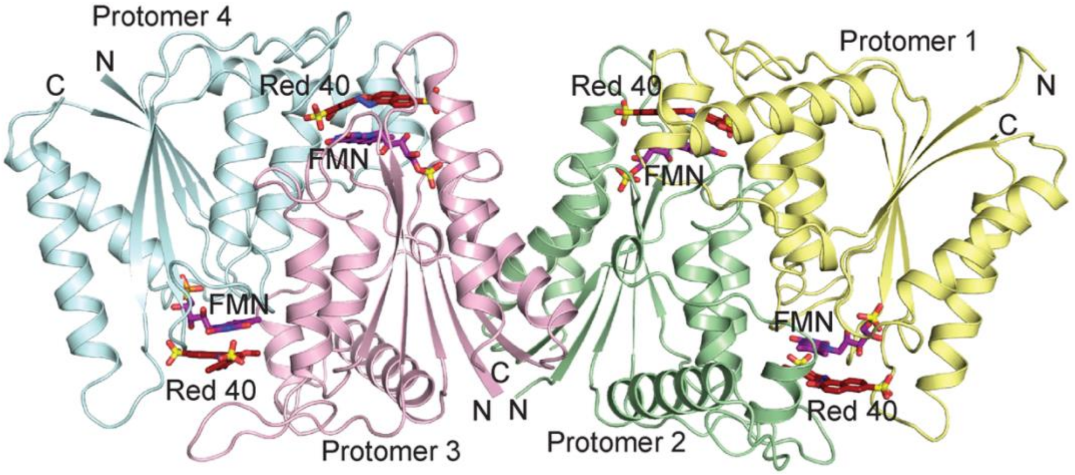

**Figure S6. Structure of AzoA bound to FMN and Red 40.** Four protomers (1-4) in the asymmetric unit of AzoA co-crystallized with Red 40 shown as in Figure 4A. FMN and Red 40 are shown as in Figure 5D.

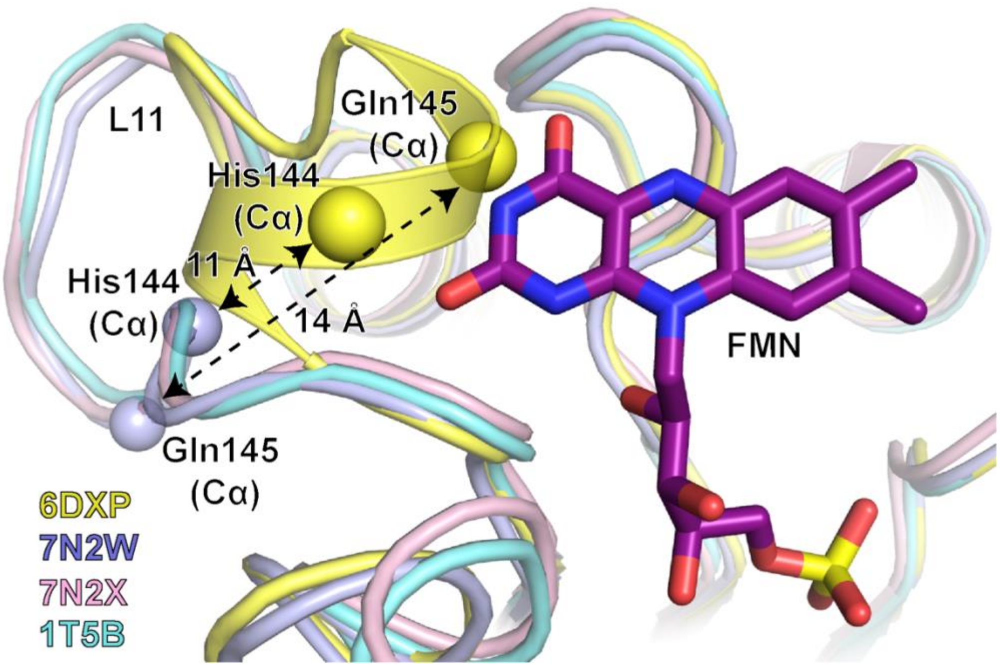

**Figure S7: Comparison of the L11 loop conformation in enzymes from Cluster 1.** Superimposed crystal structures of *K. pneumoniae* AzoA bound to Red 40 (PDB ID 7N2W; light blue), enzyme of *E. coli* AzoR (PDB 7N2X; light pink) and enzyme of *S. typhimurium* AzoR (PDB 1T5B; cyan) onto the *K. pneumoniae* AzoA holoenzyme (PDB 6DXP; yellow) highlights the pronounced change in L11 conformation in *K. pneumoniae* AzoA upon binding to substrate. Cα atoms of His144 and Gln145 from *K. pneumoniae* AzoA with or without Red 40 are shown as spheres to highlight a 11 Å and 14 Å displacement, respectively, of the L11 loop upon Red 40 binding.

### Table S1. Summary of previously characterized azoreductase-like enzymes and their clusters in Figure 1

| Species | Gene Name | Validated azoreductase activity; Reference/PMID | Cluster ID in Figure 1 |
|---|---|---|---|
| *Klebsiella pneumoniae* | AzoA | No; this paper | 1 |
| *Pseudomonas aeruginosa* | YieF | Yes; PMID:26621870 | 3 |
| *Pseudomonas aeruginosa* | ArsH | Yes; PMID:26621870 | 6 |
| *Pseudomonas aeruginosa* | MdaB | Yes; PMID:26621870 | Unlabeled two-node cluster with NP_417500.1 NADPH:quinone oxidoreductase MdaB [*Escherichia coli* str.K12 substr MG1655] |
| *Pseudomonas aeruginosa* | WrbA | No; PMID:26621870 | Unlabeled two-node cluster with WP_010877850.1 NAD(P)H:quinone oxidoreductase [*Archaeoglobus fulgidus*] |
| *Pseudomonas aeruginosa* | PA1224 | Partial; Holland, S 2017; Investigating azoreductases and NAD(P)H dependent quinone oxidoreductases in *Pseudomonas aeruginosa*, Doctor of Philosophy (PhD), Kingston University. https://ethos.bl.uk/OrderDetails.do?uin=uk.bl.ethos.733458 | 12 |
| *Pseudomonas aeruginosa* | PA1225 | No; Holland, S 2017; Investigating azoreductases and NAD(P)H dependent quinone oxidoreductases in *Pseudomonas aeruginosa*, Doctor of Philosophy (PhD), Kingston University. https://ethos.bl.uk/OrderDetails.do?uin=uk.bl.ethos.733458 | Unpaired |
| *Pseudomonas aeruginosa* | PA4975 | No; Holland, S 2017; Investigating azoreductases and NAD(P)H dependent quinone oxidoreductases in *Pseudomonas aeruginosa*, Doctor of Philosophy (PhD), Kingston University. https://ethos.bl.uk/OrderDetails.do?uin=uk.bl.ethos.733458 | 12 |
| *Bacillus subtilis* | AzoR | Yes; PMID: 16861800 | 1 |
| *Enterococcus faecalis* | AzoR | Yes; PMID: 28545445 | 1 |
| *Cereibacter sphaeroides* | AzoR | Yes; PMID: 15212802 | 1 |
| *Xenophilus azovorans* | AzoR | Yes; PMID: 20508929 | 1 |
| *Escherichia coli* | MdaB | Unknown | Unlabeled two-node cluster with NP_251270.1 hypothetical protein PA2580 [*Pseudomonas aeruginosa* PAO1] |
| *Escherichia coli* | YieF | Unknown | 3 |
| *Escherichia coli* | WrbA | Unknown | Unclustered |
| *Archaeoglobus fulgidus* | NQO | Unknown | unlabeled two-node cluster with NP_249640.1 NAD(P)H dehydrogenase [*Pseudomonas aeruginosa* PAO1] |
| *Paracoccus denitrificans* | NQO | Unknown | unclustered |
| *Sinorhizobium meliloti* | ArsH | Yes; PMID: 17673204 | 6 |

| | | | |
|---|---|---|---|
| *Desulfovibrio gigas* | | Unknown | 1 |
| *Pseudomonas luteola* | AzoR | Yes; PMID:11471684; 11101324 | 1 |
| *Pigmentiphaga kullae* | YwnB | Yes; PMID:12719939 | Unlabeled 7-sequence cluster with AAO39146.1, ADD80733.1, ANY88020.1, BAT61317.1, EKF64660.1, SUX42974.1, VEE56921.1 |
| *Geobacillus stearothermophilus IFO13737* | | Yes; PMID: 16861800 | 11 |
| *Rattus norvegicus* | NQO1 | Unknown | 12 |
| *Homo sapiens* | NQO1 | Unknown | 12 |

#### Table S2. Summary of expressed AzoA orthologs with accession codes, host and apparent molecular weight (Mol. Wt.).

| UniProt ID | Organism | Mol. Wt (kDa) |
|---|---|---|
| **P41407** | *Escherichia coli* str. K12 | 24.2 |
| **Q88IY3** | *Pseudomonas putida* KT2440 | 23.9 |
| **Q5M6G7** | *Streptococcus thermophilus* LMG18311 | 22.8 |
| **R5ZNP0** | *Streptococcus thermophilus* CAG:236 | 26 |
| **Q0TSZ6** | *Clostridium perfringens* ATCC13124 | 24.9 |
| **A6TG22** | *Klebsiella pneumoniae* subsp. pneumoniae MGH78578 | 23 |
| **A6T869** | *Klebsiella pneumoniae* subsp. pneumoniae MGH78578 | 34.8 |
| **A6TCS9** | *Klebsiella pneumoniae* subsp. pneumoniae MGH78578 | 24.2 |
| **P0AGE7** | *Escherichia coli* O157:H7 | 22.9 |

#### Table S3. The activity screen of the AzoA against menadione, AQDS (organosulfonate), various azo and quinone substrates.

| Substrate | Mean Activity (μM/min) |
|---|---|
| **Menadione** | 98.18 ± 0.33 |
| **Anthraquinone-2,6-disulfonate (AQDS)** | 22.7 ± 0.52 |
| **Balsalazide (BSZ)** | 2.98 ± 0.26 |
| **Tartrazine** | 2.43 ± 0.07 |
| **Methyl Red** | 2.26 ± 0.10 |
| **Ponceau S** | 1.78 ± 0.17 |
| **Yellow 6** | 1.44 ± 0.21 |
| **Congo Red** | 1.24 ± 0.03 |
| **Eltrombopag** | 0.96 ± 0.20 |
| **Sulfasalazine (SSZ)** | 0.45 ± 0.18 |
| **Red 40** | 0.14 ± 0.11 |
| **Phenol Blue** | -0.18 ± 0.14/ND* |
| **Benzoquinone** | -0.42 ± 0.23/ ND* |

* Not detected

### Table S4. Crystallographic data statistics and RCSB IDs

| UniProt ID | A6TCS9 | A6TCS9 | P41407 |
|---|---|---|---|
| **Bound Ligand** | FMN | FMN and Red 40 | FMN |
| **Data Collection*** | | | |
| **Source** | APS ID-31 | APS ID-31 | APS ID-31 |
| **Wavelength (Å)** | 0.98 | 0.98 | 0.98 |
| **Number of crystals** | 1 | 1 | 1 |
| **Space group** | C2 | $P2_1$ | $P42_12$ |
| **Cell dimensions** | | | |
| **a,b,c (Å)** | 133.67, 97.32, 77.53; b=90.7° | 60.41, 83.33, 87.31; b=100.3° | 94.36, 94.36, 53.1 |
| **Resolution (Å)** | 20-2.48 (2.61-2.48) | 20-2.65 (2.78-2.65) | 20-1.7 (1.73-1.7) |
| **Completeness (%)** | 98.6 (91.5) | 96.2 (81.4) | 99.9 (100.00) |
| **Total reflections** | 129402 | 84073 | 386682 |
| **Unique reflections** | 34876 | 23952 | 26943 |
| **Wilson B-factor** | 48.04 | 34.6 | 23.1 |
| **Multiplicity** | 3.7 (3.1) | 3.5 (2.5) | 14.4 (14.5) |
| **$R_{merge}$ (%)** | 10 (64.2) | 17.4 (70.0) | 5.7 (65.0) |
| **$CC_{1/2}$ (%)** | 99.6 (71.0) | 98.6 (65.3) | 99.9 (95.0) |
| **CC* (%)** | 100 (91.1) | 99.6 (88.9) | 100 (98.7) |
| **I/σ(I)** | 5.6 (1.81) | 5.4 (1.83) | 27.2 (4.3) |
| **Refinement*** | | | |
| **Reflections: work/free** | 33185 (3099)/1641 (181) | 23893 (2059)/1205 (96) | 26916 (2509)/1360 (143) |
| **$R_{work}/R_{free}$ (%)** | 23.7 (43.7)/28.5 (44.1) | 24.9 (33.8)/30.2 (33.6) | 16.5 (20.8)/19.0 (26.3) |
| **Number of TLS groups** | 0 | 22 | 6 |
| **Number of atoms** | | | |
| **Protein** | 6102 | 6003 | 1588 |
| **Ligand** | 124 | 244 | 46 |
| **Water** | 21 | 44 | 170 |
| **Average B-factors ($Å^2$)** | | | |
| **Protein** | 49.4 | 38.5 | 30.6 |
| **Ligand** | 39.1 | 44.7 | 31.4 |
| **Water** | 28.5 | 33.9 | 35.3 |
| **RMSD** | | | |
| **Bond lengths (Å)** | 0.012 | 0.01 | 0.007 |
| **Bond angles (°)** | 1.58 | 1.35 | 1 |
| **Molprobity †** | | | |
| **Favored** | 98.2% (746 aa) | 99.6% (731 aa) | 99.6% (202 aa) |
| **Allowed** | 99% (751 aa) | 100% (746 aa) | 100% (203 aa) |
| **Outliers** | 1% (9 aa) | none | none |
| **Clash score** | 3.62 | 6.45 | $100^{th}$ percentile |
| **MolProbity score** | 1.76 | 1.96 | $100^{th}$ percentile |
| **PDB ID** | **6DXP** | **7N2W** | **7N2X** |

*Statistics calculated using PHENIX.[8]
†Calculated with the program MolProbity.[10]

### References

(1) Gileadi, O.; Burgess-Brown, N. A.; Colebrook, S. M.; Berridge, G.; Savitsky, P.; Smee, C. E.; Loppnau, P.; Johansson, C.; Salah, E.; Pantic, N. H. High throughput production of recombinant human proteins for crystallography. *Methods Mol Biol* **2008**, *426*, 221–246. DOI: 10.1007/978-1-60327-058-8_14.

(2) Studier, F. W. Protein production by auto-induction in high-density shaking cultures. *Protein Expression and Purification* **2005**, *41* (1), 207–234. DOI: 10.1016/j.pep.2005.01.016 (accessed 2021-01-08T12:02:47).

(3) Battye, T. G.; Kontogiannis, L.; Johnson, O.; Powell, H. R.; Leslie, A. G. iMOSFLM: a new graphical interface for diffraction-image processing with MOSFLM. *Acta Cryst* **2011**, *D67* (Pt 4), 271–281. DOI: 10.1107/S0907444910048675.

(4) Evans, P. Scaling and assessment of data quality. *Acta Cryst* **2006**, *D62* (Pt 1), 72–82. DOI: 10.1107/s0907444905036693 From NLM.

(5) McCoy, A. J.; Grosse-Kunstleve, R. W.; Adams, P. D.; Winn, M. D.; Storoni, L. C.; Read, R. J. Phaser crystallographic software. *J Appl Crystallogr* **2007**, *40*, 658–674. DOI: 10.1107/s0021889807021206.

(6) Ito, K.; Nakanishi, M.; Lee, W. C.; Sasaki, H.; Zenno, S.; Saigo, K.; Kitade, Y.; Tanokura, M. Three-dimensional structure of AzoR from *Escherichia coli*. An oxidoreductase conserved in microorganisms. *J Biol Chem* **2006**, *281* (29), 20567–20576.

(7) Murshudov, G. N.; Skubak, P.; Lebedev, A. A.; Pannu, N. S.; Steiner, R. A.; Nicholls, R. A. REFMAC5 for the refinement of macromolecular crystal structures. *Acta Crystallogr* **2011**, *D67*, 355–367. DOI: 10.1107/s0907444911001314.

(8) Liebschner, D.; Afonine, P. V.; Baker, M. L.; Bunkóczi, G.; Chen, V. B.; Croll, T. I.; Hintze, B.; Hung, L. W.; Jain, S.; McCoy, A. J.; et al. Macromolecular structure determination using X-rays, neutrons and electrons: recent developments in PHENIX. *Acta Crystallogr* **2019**, *D75* (10), 861–877. DOI: 10.1107/s2059798319011471 From NLM.

(9) Emsley, P.; Lohkamp, B.; Scott, W. G.; Cowtan, K. Features and development of Coot. *Acta Cryst* **2010**, *D66* (Pt 4), 486–501, Research Support, Non-U.S. Gov't. DOI: 10.1107/S0907444910007493.

(10) Chen, V. B.; Arendall, W. B., 3rd; Headd, J. J.; Keedy, D. A.; Immormino, R. M.; Kapral, G. J.; Murray, L. W.; Richardson, J. S.; Richardson, D. C. MolProbity: all-atom structure validation for macromolecular crystallography. *Acta Crystallogr D* **2010**, *66* (Pt 1), 12–21.
